# Identification of cell types associated with 14 brain phenotypes from more than 10 million single cells

**DOI:** 10.64898/2025.12.05.692533

**Authors:** Tanya N Phung, Sara L Seoane, Wei-Ping Li, Rachel Brouwer, Danielle Posthuma

**Affiliations:** Department of Complex Trait Genetics, Center for Neurogenomics and Cognitive Research, Vrije Universiteit Amsterdam, Netherlands, 1081HV; Department of Child and Adolescent Psychology and Psychiatry, section Complex Trait Genetics, VU University Medical Center, Amsterdam, The Netherlands, 1081HV

## Abstract

Genome-wide association studies (GWAS) have yielded unprecedented insight into the genetic variants associated with many human traits. However, translating this insight into knowledge about causal biological mechanisms remains challenging. One promising recent strategy is to link GWAS-associated genes to information on expression levels of those genes in specific cell types. This strategy allows for the generation of specific, testable hypotheses about which cell types are important for a trait which can then be investigated in functional lab experiments for actual relevance. The success of this strategy strongly depends on the quality and systematic analysis of available single-cell RNAseq datasets. Here we present a comprehensive database of 388 datasets derived from 36 studies spanning different regions of the brain across developmental timepoints. Using this database, we tested for the presence of cell type enrichment in genes associated with 14 traits. We confirmed previous findings such as the association between microglia and Alzheimer’s disease in the entorhinal cortex. In addition, we found novel evidence for the involvement of specific cell types in disease, such as astrocytes being implicated in alcohol-related phenotypes or neuronal cell types at the prenatal stage in ADHD. Our database has been incorporated to FUMA, a publicly available and widely used tool for post-GWAS functional annotation analyses. Our work provides an approach to facilitate the prioritization of cell types in specific brain regions and/or developmental stages, further informing the design of follow-up experiments.

## Introduction

The past twenty years of Genome wide association studies (GWAS) have resulted in the discovery of thousands of genetic variants associated with complex traits (Cerezo et al., 2025). However, the polygenic nature of complex traits has made it difficult to translate these discoveries into mechanistic insights about causal mechanisms. One strategy that is often employed to gain biological insights is to test if trait-associated variants are enriched in biological functions or pathways. Advances in single-cell technology makes it now possible to sequence single cells on the orders of thousands to millions of cells (hereafter we used single-cell to include both single-cell and single-nuclei RNAseq datasets). Recent large-scale examples include the Human Brain Cell Atlas in which about 3 million cells obtained from 3 male donors across 105 human brain regions were sequenced (Siletti et al., 2023). PsychENCODE Consortium sequenced multi-omics single-cells from >2.8 million cells from brain tissue of the prefrontal cortex of 388 individuals (Emani et al., 2024). Leveraging these large-scale single-cell resources, recent studies tested if GWAS variants from brain phenotypes converge in specific cell types. This approach has generated concrete, testable hypotheses for brain-related traits. For example, Ma et al. 2023 utilized a single-cell RNA-seq dataset from the human entorhinal cortex to identify significant associations of microglia and oligodendrocyte precursor cells (OPCs) to Alzheimer’s disease, which led to the specific hypothesis that pathways relevant to brain development and synaptic transmitters could contribute to the pathogenicity of Alzheimer’s disease (Ma et al., 2023). Duncan et al. 2025 identified specific neuronal subtypes associated with schizophrenia in the hypothalamus leading to the hypothesis that these neuronal subtypes may explain the reduction of hippocampal volume in schizophrenia patients (Duncan et al., 2025). Overall, integrating GWAS signals and single-cell RNAseq data has greatly expanded our understanding of the biological mechanisms underlying brain-related traits. Further, identifying associated cell types can lead to new directions in therapeutic development, as in the case of targeting microglia in Alzheimer’s disease (Zhang et al., 2021).

Up to now, GWAS to cell type prioritization studies have used only a few (<5) gene expression datasets (Supplementary Table 1). This is limiting since these single-cell brain datasets typically include brain tissue from a small number of donors, thereby ignoring individual differences and in particular potential sex-specific effects (as in the case of 3 male donors in Siletti et al., 2023), or are limited to a specific brain region or developmental stage. Even in studies that aimed to examine cell types from different brain regions, they were limited to mouse brain (Bryois et al., 2020).

Due to these limitations and due to the speed with which novel single-cell RNAseq resources have become available, focusing on the human brain, we set out to 1) harmonize high quality single-cell RNA-seq resources, and 2) use this central database to test for cell type enrichment across 14 traits. Since we included spatial and temporal gene expression information, we also tested cell type enrichment across different brain regions across different developmental times. Because conducting analyses to identify trait-associated cell types from GWAS variants and single-cell RNAseq datasets often hinges on having the single-cell RNAseq datasets systematically processed, we incorporated these processed datasets into an online repository (FUMA) to facilitate rapid utilization of these resources by the scientific community.

## Methods

### Curation and processing of single-cell RNAseq datasets from human brain

We curated publicly available single-cell RNAseq datasets from the cellxgene resource (link: https://cellxgene.cziscience.com/collections) in three batches, performed on May 29, 2024, February 9, 2025, and September 2 2025. We applied a filter to include the single-cell RNAseq datasets that were sequenced from the human brain of un-diseased individuals (Supplementary Note 1). Ultimately, we proceeded with 36 datasets for further processing (Table 1). Details about included studies and exclusion criteria can be found in Supplementary Table 2 and Supplementary Note 1.

**Table 1.** Single-cell RNAseq datasets curated.

| Authors/Year | Brain region | Developmental stage | Number of cells (approximate) |
| --- | --- | --- | --- |
| (Smith et al., 2021) | neocortex | midgestational | ~120k |
|  |  | infant | ~50k |
| (Jäkel et al., 2019) | white matter | adult | ~7k |
| (Leng et al., 2021) | entorhinal cortex | adult | ~10k |
|  | superior frontal gyrus |  | ~12k |
| (Aldinger et al., 2021) | cerebellum | 1st and 2nd trimester | ~69k |
| (Bhaduri et al., 2021) | neocortex | 2nd trimester | ~450k |
| (Bakken et al., 2021) | primary motor cortex | adult | ~20k |
| (Kamath et al., 2022) | substantia nigra | adult | ~370k |
| (Otero-Garcia et al., 2022) | prefrontal cortex | adult | ~58k |
| (S. Ma et al., 2022) | dorsolateral prefrontal cortex | adult | ~170k |
| (Gabitto et al., 2024) | middle temporal gyrus | adult | ~160k |
| (Seeker et al., 2023) | white matter | adult | ~45k |
| (Gittings et al., 2023) | frontal and occipital cortex | adult | ~100k |
| (Jorstad, Song, et al., 2023) | middle temporal gyrus | adult | ~150k |
| (Siletti et al., 2023) | 10 brain regions | adult | ~3million |
| (Zhu et al., 2023) | neocortex | early mid gestation fetal, late mid gestation fetal, infancy, childhood, adolescence, and adulthood | ~50k |
| (Jorstad, Close, et al., 2023) | neocortex | adult | ~1million |
| (Velmeshev et al., 2023) | cortex and ganglionic eminence | 2nd trimester, 3rd trimester, 0-1 years, 1-2 years, 2-4 years, 4-10 years, 10-20 years, and Adult | ~700k |
| (Sepp et al., 2024) | cerebellum | 1st trimester, 2nd trimester, newborn, infant, toddler, and adult | ~160k |
| (Phan et al., 2024) | caudate nucleus and putamen | adult | ~45k |
| Nascimento | entorhinal cortex and ganglionic eminence | fetal, infant, toddler, teen, adult | ~150k |
| TorresFlores | cerebellum | child | ~1k |
| (Wang et al., 2025) | neocortex | 1st trimester, 2nd trimester, 3rd trimester, infancy, adolescence | ~230k |
| (Braun et al., 2023) | cerebellum, diencephalon, forebrain, head, hindbrain, medulla, midbrain, pons,<br>telencephalon | 1st trimester | ~1.6million |
| (Rexach et al., 2024) | primary visual cortex,<br>insular cortex,<br>Brodmann (1909) area<br>4 | adult | ~100k |
| (Dharshini et al., 2024) | Brodmann (1909) area<br>9, Brodmann (1909)<br>area 7, Brodmann<br>(1909) area 17 | adult | ~100k |
| (N. M. et al., 2024) | dorsal motor nucleus<br>of vagus nerve,<br>primary visual cortex,<br>prefrontal cortex,<br>primary motor cortex,<br>medial globus pallidus | adult | ~500k |
| (Clarence et al., 2025) | caudate nucleus,<br>hippocampal<br>formation, dorsolateral<br>prefrontal cortex,<br>anterior cingulate<br>cortex | infant stage,<br>adolescence, adult | ~100k |
| (Tadross et al., 2025) | hypothalamus | adult | ~430k |
| (Pan et al., 2024) | prefrontal cortex | adult | ~30k |
| (Xu et al., 2023) | hippocampus | adult | ~650k |
| (Johansen et al.,<br>2023) | Neocortex | adult | ~380 |
| (Han et al., 2020) | adult cerebellum, fetal<br>brain | fetal and adult | ~20k |
| (Cao et al., 2020) | cerebellum and<br>telencephalon | prenatal | ~700k |
| Linnarson | brain meninx,<br>forebrain meninges,<br>meninx | 1st trimester | ~150k |
| Yang | dorsolateral prefrontal<br>cortex | adult | ~20k |
| (Lee et al., 2024) | dorsolateral prefrontal<br>cortex | adult | ~120k |

### Subset single-cell RNAseq studies to brain regions and developmental timepoints

The downloaded single-cell RNAseq files from the included studies typically contain gene expression data for all cells within a study, which in some cases stem from different brain regions or different developmental stages. We created subsets of specific brain regions and developmental stages using the metadata available from the downloaded data (for more details, refer to Supplementary Note 1). This procedure resulted in 388 datasets (Supplementary Table 3). From here on, ‘parent study’ refers to the original 36 studies and ‘datasets’ refers to the 388 datasets that were derived from the original 36 studies when separating into brain regions and developmental stages. For interpretability, we further grouped the 36 brain regions into 7 groups: broad embryological divisions, developmental derivatives of the embryological divisions, cortical types based on laminar structure, non-neocortical, lobar subdivisions of neocortex based on anatomy and function, fine-grained cortical areas with specific cognitive/sensory/motor functions, and non-laminar brain structures (Supplementary Table 4). The curated single-cell brain datasets span two developmental stages: prenatal (defined as pre-birth) and postnatal (defined as post-birth) (Supplementary Table 5). Note that in some prenatal datasets, there was no additional resolution other than ‘brain’. We included these datasets in our analyses and considered them as a separate group.

### Quality control

For each of the 388 single-cell RNAseq datasets, cells were kept if they met the following filtering criteria: cells with at least 200 detected genes and cells where the mitochondrial percentage was less than ten percent. Additionally, we only kept genes that were expressed in at least three cells. For consistency, we converted gene symbols to Ensemble ID in all datasets.

### Assignment of author-labelled cell types to broad categories

Supplementary Table 6 lists all 283 unique cell type labels from the 36 single-cell RNAseq datasets in this study. We observed that the author-labeled cell types tend to fall into four main categories: progenitor, neuron, glia, and other (which includes support cell types such as epithelial cells and muscle cells, immune cell types and ambiguous labels such as unknown or miscellaneous). We first assigned each of the 283 unique cell type labels to a broad category (progenitor, neuron, glia, and other) (Supplementary Table 6). Where it was possible to do so, we assigned author-defined cell types to more specific labels. The progenitor label was further divided into neuronal progenitor (to indicate the progenitor cells that are most likely to differentiate into neuronal cell types), radial glial cell, and glial progenitor; the neuron label was further divided into generic neuron (to include the labels that are generic and cannot be classified further, such as neuron or central nervous system neuron), excitatory neuron, inhibitory/interneuron, and specialized neuron (to include the specialized labels such as hippocampal granule cell); the glia label was further divided into oligodendrocyte precursor, oligodendrocyte, astrocyte, and microglia.

### Selection of GWAS summary statistics

To identify well-powered, publicly available GWAS, we downloaded the metadata file from GWAS catalog (https://www.ebi.ac.uk/gwas/api/search/downloads/studies/v1.0.3.1; access date: January 10 2025). Out of 126,995 available records, 85,932 records had full summary statistics available. We then selected brain-related phenotypes where the total GWAS sample size was at least 500,000 individuals in the European population, which resulted in 27 phenotypes (Supplementary Table 7, ID 1 to 27). For binary phenotypes, we selected on the total sample size defined as the sum of number of cases and number of controls. Additionally, we added seven brain-related phenotypes from the Psychiatric Genomics Consortium (Supplementary Table 7, ID 28 to 34). We also selected body mass index (BMI) from (Yengo et al., 2018). Even though BMI is a metabolic trait, previous research has suggested the associations between body mass index and the brain (Fusco et al., 2025). The remaining 35 phenotypes contained similar phenotypes, of which we selected one phenotype with the highest sample size overall for quantitative traits and the highest proportion of cases for binary traits (Supplementary Note 2). In summary, we selected 14 traits that are representative of cognitive, psychiatric and neurological disorders (Table 2, Supplementary Table 8).

**Table 2.** Phenotypes tested. The symbol ∼ is used to denote the approximation of sample size from averaging the sample size column from the summary statistics

| Phenotype<br>(Abbreviation) | Domain | Type | Number<br>of cases | Number of<br>controls | Total<br>sample<br>size | Authors/Year |
| --- | --- | --- | --- | --- | --- | --- |
| Cognitive processing<br>speed | cognitive | quantitative | N/A | N/A | 2,266,733 | (Li et al., 2024) |
| Well-being | psychiatric | quantitative | N/A | N/A | 2,083,151 | (Baselmans et al., 2019) |
| Smoking initiation | psychiatric | binary | 312584 | 320218 | 1,232,091 | (Liu et al., 2019a) |
| Depression | psychiatric | binary | 525197 | 3362335 | 3887532 | (Adams et al., 2025) |
| Schizophrenia (SCZ) | psychiatric | binary | 55193 | 74132 | 129325 | (Trubetskoy et al., 2022) |
| Alcohol consumption | psychiatric | quantitative | N/A | N/A | 941280 | (Liu et al., 2019b) |
| Neuroticism | psychiatric | quantitative | N/A | N/A | 682688 | (Gupta et al., 2024) |
| ADHD | psychiatric | binary | 38691 | 186843 | 225534 | (Demontis et al., 2023) |
| Bipolar Disorder | psychiatric | binary | 59287 | 781022 | 840309 | (O'Connell et al., 2025) |
| Parkinson's disease (PD) | neurological | binary | ~17675 | ~239152 | ~256827 | (Nalls et al., 2019) |
| Migraine | neurological | binary | 28852 | 525717 | 554569 | (Choquet et al., 2021) |
| Alzheimer's disease (AD) | neurological | binary | 156135 | 2049717 | 2205852 | (Uffelmann et al., 2025) |
| Insomnia | neurological | binary | 109548 | 277440 | 386988 | (Watanabe et al., 2022) |
| Body mass index (BMI) | metabolic | quantitative | N/A | N/A | ~700000 | (Yengo et al., 2018) |

### Predict relevant cell types for each GWAS

To identify which cell types are associated with a trait, we modified the strategy described in (Watanabe et al., 2019) (Figure 1). Given that cell types are likely to be similar across our datasets, by design, either because they stem from the same parent study or because the studies captured similar cell types, we expect multiple cell types to be associated with each trait. In case of cell types from different parent studies, this is not always easy to see because cell type labels are not always consistent. To identify the number of ‘independent’ cell types, we implemented a three step conditional procedure. For each of the 388 datasets, for each cell type defined by the studies’ authors, we computed mean in expression across all cells with the same cell type label. Then, for each GWAS, we applied MAGMA gene property analysis (step 1 of the FUMA Cell type procedure) for each dataset. Average expression across cell types was added as an additional covariate. We interpret the cell types that are significantly associated with a phenotype post MAGMA gene property analyses (step 1) as putatively relevant cell types. Given that two or more datasets originate from the same ‘parent study’ and therefore contain a similar set of cell types, these analyses are not independent. Thus, while in Watanabe et al. 2019, Bonferroni correction was based on the total number of cell types, we modified the Bonferroni correction to be based on the number of unique cell type labels across datasets per brain region. Subsequently, significant cell types from step 1 were selected based on Bonferroni corrected p value < 0.05 where the number of tests is defined as the total unique cell types across all datasets within a brain region (see Supplementary Table 9 for the number of unique cell types per region). Note that we analyzed datasets within prenatal and postnatal timepoints separately.

**Figure 1.**
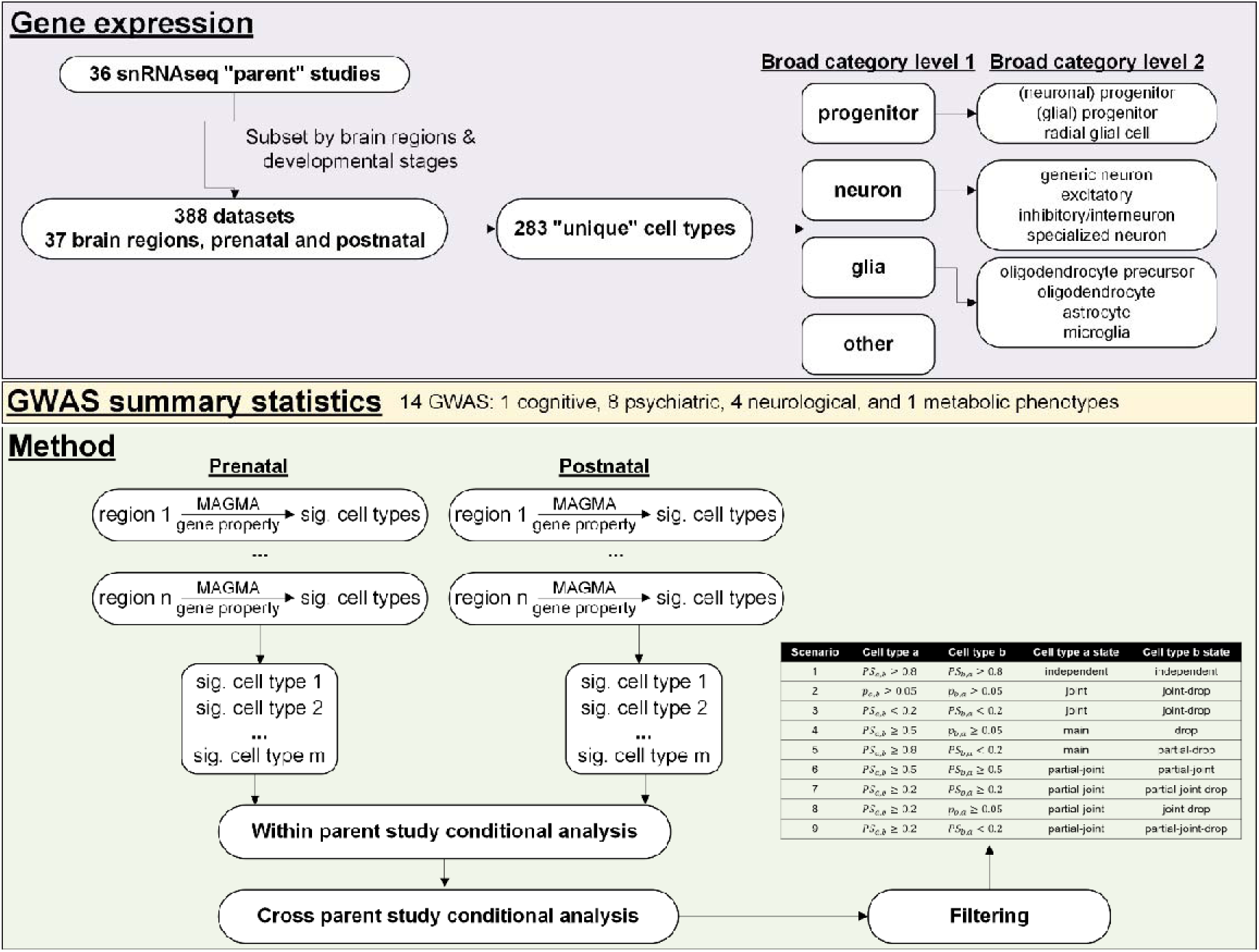
Overview of study design.

We then merged significant cell types post step 1 across all brain regions and implemented a within ‘parent study’ conditional analysis (step 2 of the FUMA Cell type workflow): the rationale of the step 2 in Watanabe et al. 2019 is to perform conditional analyses on cell types within the same dataset. In our case, because two or more datasets within the same region and cross brain regions could originate from the same parent study, we modified the step 2 strategy to perform conditional analyses on all cell types within the same ‘parent study’. Cell types were selected for further analysis in step 3 according to the criteria described in Watanabe et al., 2019, which implements a forward selection procedure for a pair of cell types. The selected cell types are interpreted to be independently associated to the phenotype.

Finally, we implemented cross dataset conditional analysis (step 3 of the FUMA Cell type workflow): here we performed cross ‘parent study’ conditional analyses. We implemented the same selection criteria as in step 2 of the workflow. This step results in clusters of cell types that are associated with the phenotype where each cluster contains a set of cell types that should, in theory, be the same underlying cell type (leaves in the illustrative dendrogram plots in Supplementary Note 3). We interpret the remaining cell types after step 3 as representatives for a group of transcriptionally similar cell types. We used the main cell type in the cross ‘parent study’ conditional analyses to name each cluster (nodes in the illustrative dendrogram plots in Supplementary Note 3). For an overview of cell types associated with phenotypes, we mapped the main cell types to a broad category (Supplementary Table 6).

### Code and data availability

All of the datasets processed here have been added to FUMA Cell Type (version 1.8.2). Codes used to process the scRNAseq data: https://github.com/tanyaphung/scrnaseq_viewer. Codes used for the project: https://github.com/tanyaphung/brain_celltypes. Codes used for the modified FUMA Celltype workflow: https://github.com/tanyaphung/FUMA_Celltype_cmd.

## Results

### Overview of single-nuclei expression datasets

In the 36 processed single-cell RNAseq datasets, single-cell RNAseq data in certain brain regions are only available either prenatally or postnatally (Figure 2). The unavailability of single-cell RNAseq for a brain region could be due to inability to define brain regions prenatally (in other words, the brain at the first trimester might not have further developed to specialized regions) and some regions such as transient structure of the forebrain do not exist postnatally. The lack of snRNA-seq datasets for some regions reflects a combination of biological and methodological factors. During early prenatal development (e.g., first trimester), many adult-like anatomical subdivisions have not yet emerged and therefore cannot be reliably defined or dissected (e.g., Polioudakis et al., 2019)), and other regions (such as transient forebrain structures) are developmentally restricted and disappear postnatally (Judaš et al., 2013).

**Figure 2.**
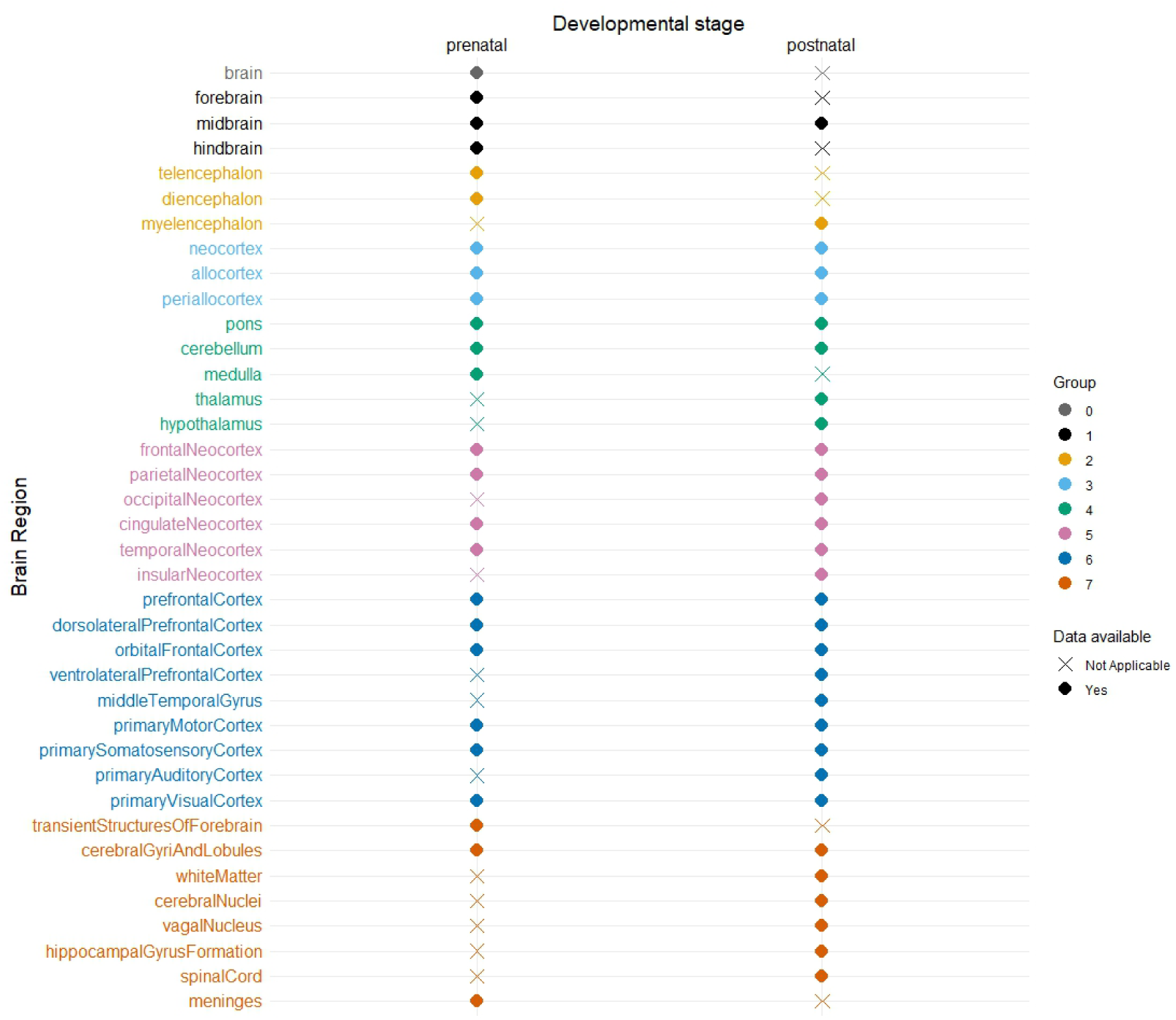
Availability of single-cell RNAseq data stratified by brain regions and developmental stages. Availability of single-cell RNAseq data for each brain region is shown: filled circle denotes single-cell RNAseq data are available for the region and an ‘x’ otherwise. Brain regions are color-coded corresponding to the 7 groups (Supplementary Table 4).

To obtain an overview of which categories of cell types were annotated in the curated 36 single-cell RNAseq datasets, we manually assigned the cell type label from each study to broad categories (Methods, Supplementary Table 6). We observed the presence of progenitor, neuronal, and glial cell types in these curated single-cell RNAseq datasets both prenatally and postnatally (Supplementary Figure 1).

Results from steps 1-3 for each of the phenotypes separately can be found in Supplemental Tables 10-37. Below, we discuss overall patterns at the broad cell type level or collapsing across brain regions.

### Cell types associated with brain phenotypes in different brain regions

Collapsing the cell types into broad categories, we observed that similar cell labels are putatively associated with studied phenotypes across all brain regions and developmental stages. A notable exception is of Parkinson’s disease (group 1: midbrain and group 4: cerebellum and medulla) and migraine (group 2: diencephalon, group 3: neocortex, group 5: frontal neocortex, parietal neocortex, cingulate neocortex, and temporal neocortex, and group 6: prefrontal cortex, dorsolateral prefrontal cortex, and primary visual cortex (Figure 3, prenatal). While it is biologically plausible that similar cell types are affected in multiple brain regions, the conditional analysis strategy allows us to identify the cell types that are most likely to be independently associated. Focusing on cell types that remain significant post conditional analyses (red squares in Figure 3), we can identify specific relevant brain regions that could be prioritized for follow-up functional analyses. We observed the association between cell types in the entorhinal cortex (brain region periallocortex) with Alzheimer’s disease and between the midbrain and Parkinson’s disease, confirming previous findings (Ma et al., 2023). Additionally, we observed new findings such as an association between cerebellar cell types and depression prenatally or between cell types in the hypothalamus and Alzheimer’s disease which are associations in brain regions that have not been selected previously in analyses linking GWAS to cell types.

**Figure 3.**
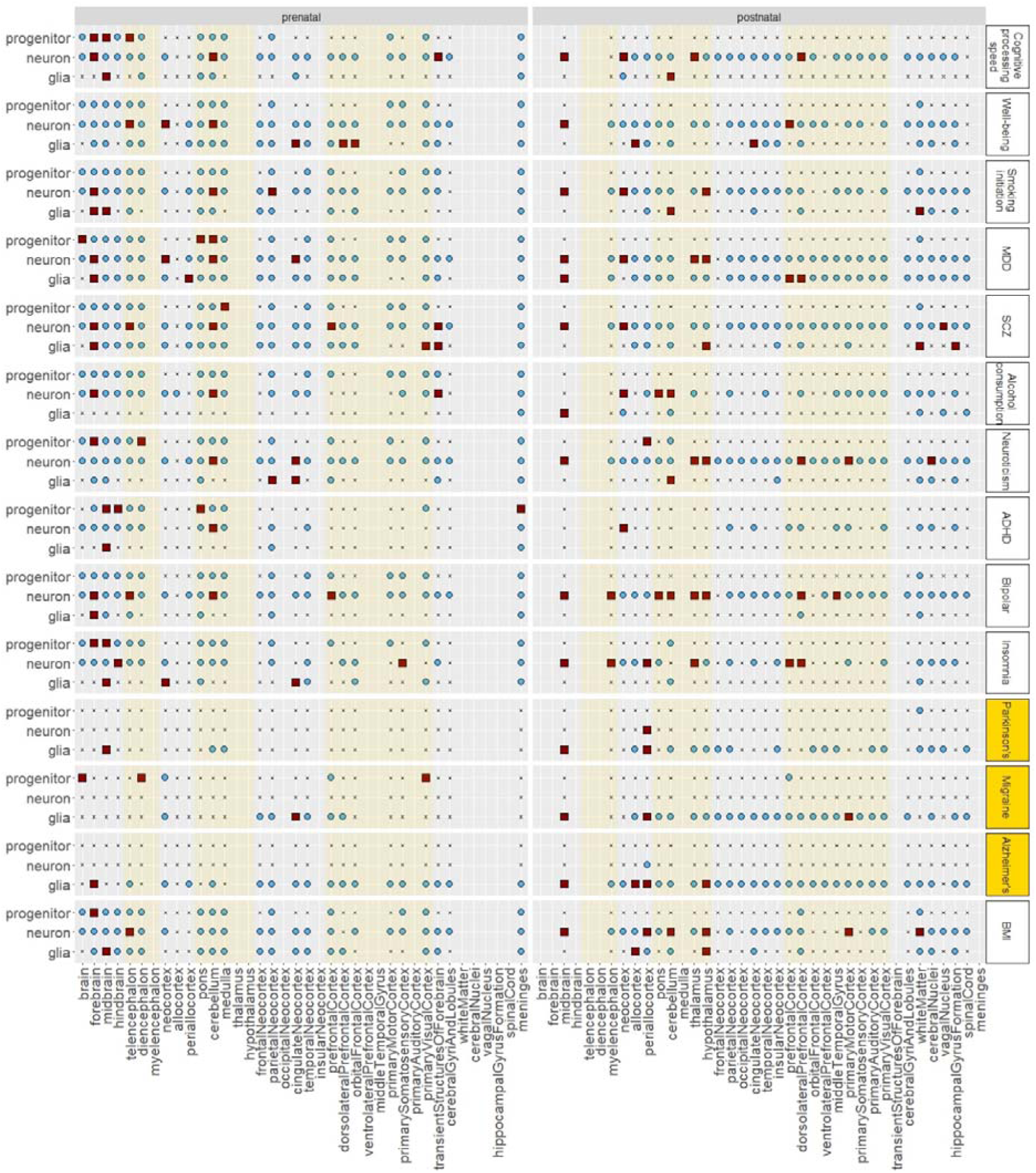
Overview of cell types associated with brain phenotypes stratified by regions. Broad category of cell type (level 1) (y-axis) that is significantly associated with phenotypes (per row) stratified by brain regions (x-axis). The category ‘other’ is not included in this plot. Blue circle denotes significantly associated in at least one cell type in this category post step 1 (MAGMA gene property). Red squares denote at least one independently associated cell type post step 3 (see Methods). The symbol ‘x’ indicates that there is no significantly associated cell type whereas an empty column indicates that there is no single-cell RNAseq data for the brain region. The light yellow boxes are used to separate out the regions. The neurological phenotypes are labelled in yellow, others in white.

### Putatively associated cell types differ between phenotypes and across developmental timepoints

To examine if there are differences in associated cell types between different types of phenotypes and developmental timepoints, we collapsed the brain regions in the next analysis. We observed differences in putatively associated cell types when comparing ‘cognitive and psychiatric’ and ‘neurological’ phenotypes. Progenitor, neuronal, and glial cell types are associated with cognitive and psychiatric phenotypes and BMI while for the three neurological phenotypes in this study, only glial (and to some extent neuronal) cell types are associated (Figure 4).

**Figure 4.**
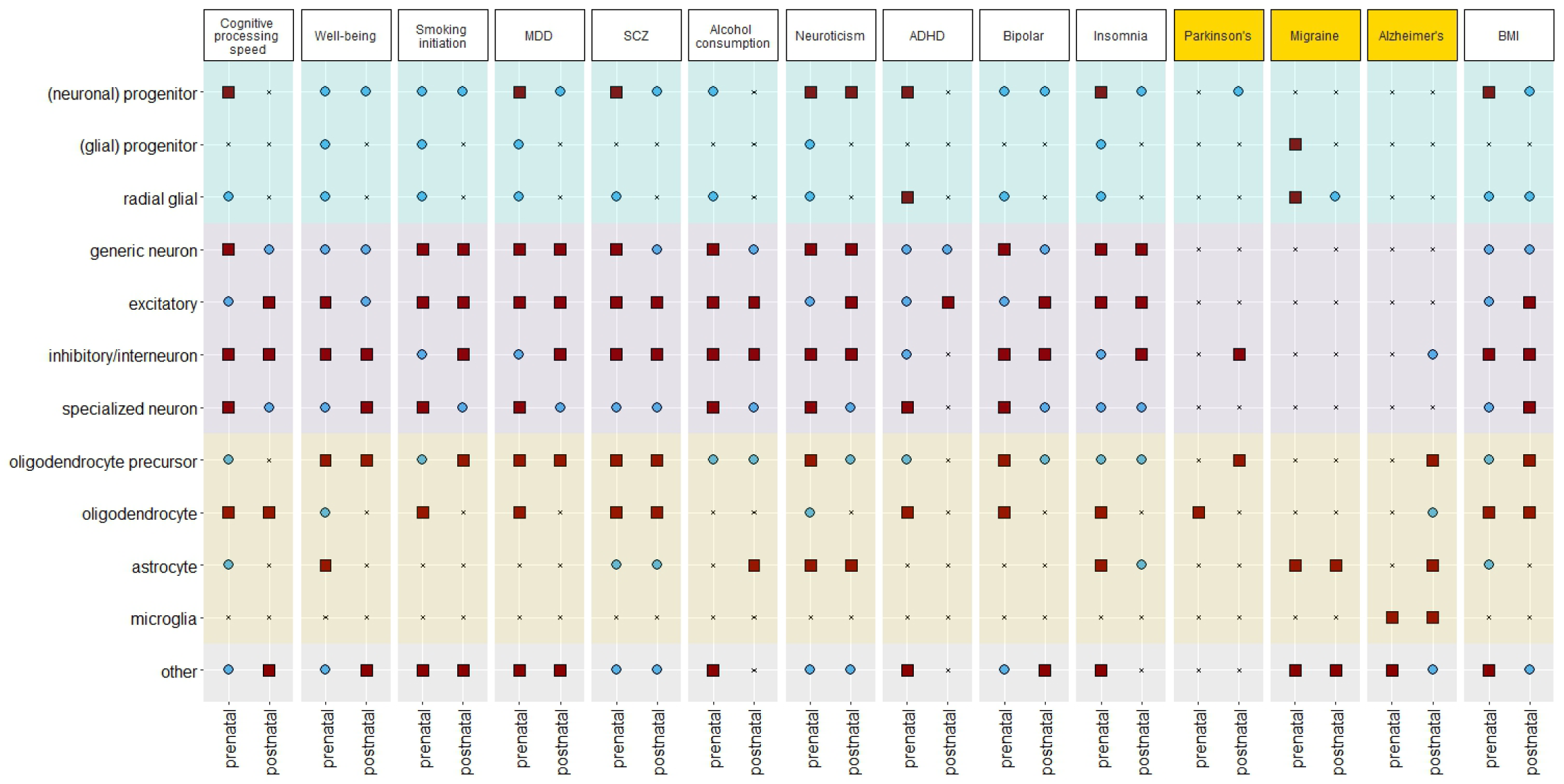
Overview of cell types associated with brain phenotypes. Broad category of cell type (level 2) (y-axis) that is significantly associated with phenotypes. In this plot, the information on specific brain region is not taken into account. Circle denotes at least one significantly associated cell types in this brain region post step 1 (MAGMA gene property). Squares denote independently associated cell types post step 3 (see Methods). Broad categories of cell type level 1 are shaded in blue for progenitor, in purple for neuron, in yellow for glia, and gray for other. The symbol ‘x’ indicates that there is no significantly associated cell type. The neurological phenotypes are labelled in yellow, others in white.

We observed specific temporal differences (Figure 4). For example, significant association with astrocyte is observed only prenatally in cognitive processing speed, well-being, and BMI, only postnatally in alcohol consumption and AD, and across both stages in schizophrenia, neuroticism, insomnia, and migraine. For Parkinson’s disease and Alzheimer’s disease, only glial cell types are associated prenatally while both neurons (specifically inhibitory/interneuron) and glial cell types are associated postnatally.

We found more categories of cell types that are associated with ADHD prenatally as compared to postnatally: only neuronal cell types postnatally but all broad categories are associated prenatally. The reverse pattern is observed in AD: only glial cell types prenatally but both neuronal and glial cell types postnatally.

### The role of non-neuronal cell types in brain phenotypes

While we confirmed the association between neuronal cell types and psychiatric phenotypes (Bryois et al., 2020; Duncan et al., 2025; Skene et al., 2018; Yao et al., 2025), here we also reported on the role of non-neuronal cell types in brain phenotypes. Many phenotypes in this study are linked to oligodendrocyte precursors, oligodendrocyte, and to a certain extent, astrocyte (Figure 4). Only migraine is not associated with either oligodendrocyte or oligodendrocyte precursors but is associated with astrocyte both prenatally and postnatally. Alcohol consumption is associated with oligodendrocyte precursor but not with oligodendrocyte. Interestingly, microglia are only associated with AD and not with any other phenotypes in this study. In addition to glial cell types, we also found that progenitor cell types, specifically progenitors that will most likely differentiate to neurons, are associated with the studied phenotypes.

## Discussion

In this study, we systematically processed publicly available single-cell RNAseq data that were sequenced from multiple regions of the non-diseased human brain at different developmental timepoints. We modified an existing approach to link GWAS findings and single-cell RNAseq data to account for the complexity created by analyzing data from different brain regions that are highly correlated. We applied this framework to gain insights into the general patterns of putatively associated cell types in 14 brain-related traits. We confirmed previous findings but also identified new associations that could be further prioritized in follow-up functional experiments.

It is thought that different brain regions are important for different brain-related phenotypes. There has been some support for this such as the hippocampus in Alzheimer’s disease (Rao et al., 2022). Recently, Yao et al. 2025 found specific brain regions (amygdala, hippocampal body, and prefrontal cortex) to be associated with schizophrenia (Yao et al., 2025). We hypothesized that we could leverage the availability of single-cell RNAseq data that were sequenced from multiple brain regions to pinpoint specific areas that are associated with brain-related traits. In contrast to our hypothesis, we found no clear distinction in putatively associated cell types across different regions of the brain, except for Parkinson’s disease and migraine at the prenatal developmental timepoint. These results could suggest the involvement of cell types from different brain regions to be important in these phenotypes. Or it could be due to limitations of the framework. It is plausible that cell types from different brain regions are highly correlated at the expression level and current methods are unable to tease out if specific regions are associated with the phenotypes (Jorstad, Song, et al., 2023). Nonetheless, leveraging the conditional analysis strategy, we reasoned that the cell types that remain significant should be prioritized in follow-up analyses. Using this rationale, we confirmed previous results such as the association between entorhinal cortex (annotated as periallocortex in this study) and Alzheimer’s disease and between the midbrain and Parkinson’s disease. We additionally prioritized specific brain regions that are involved with the phenotypes that were not known previously such as cerebellum and depression.

Our detailed systematic analysis makes it possible to identify broad differences in putatively associated cell types temporally, between different types of traits, and highlight the contribution of non-neuronal cell types in brain-related traits. Differences in the relevance of developmental versus adult cell types could be further explored to better understand the biological mechanisms throughout development. We observed clear distinction of associated cell types between Parkinson’s disease, migraine, and Alzheimer’s disease which are ‘neurological’ phenotypes versus the remaining 11 phenotypes which are cognitive and psychological phenotypes. Studies that leverage GWAS findings and single-cell RNAseq datasets to identify relevant cell types in brain-related traits often result in the prioritization of neuronal cell types (Supplementary Table 1). In addition to confirming the association with neuronal cell types, we expand the role of progenitor and glial cell types in being relevant to brain-related phenotypes.

In summary, while methodology to link GWAS findings and single-cell RNAseq datasets has proven successful in yielding insights into the biological mechanisms of polygenic phenotypes, in brain-related phenotype specifically, relying on one to two single-cell RNAseq data does not capture spatial or temporal variation. By systematically creating a single-cell RNAseq database that are publicly available from different brain regions at different developmental timepoint, we could prioritize specific brain regions and broad categories of cell types that are relevant for brain phenotypes. This knowledge can then be used in follow-up analyses to hopefully pinpoint the specific relevant genes and pathways. For example, a potential follow-up experiment based on these results could be to leverage the observation that astrocytes are associated with alcohol related disorders: one could create iPSC-cells collected from individuals with and without alcohol use disorders, differentiate them to astrocytes and study if there are genes or proteins that are differentially expressed.

To identify associated cell types from linking GWAS results and single-cell RNAseq datasets, we applied one previously developed method. Ongoing work by us and others have shown that the associated cell types that are identified by different methods do not always concur (Li et al., 2025, also work in preparation). While it might be most optimal to implement more than one method, due to the large-scale datasets, it was not computational feasible. However, we focus our interpretation on broad categories of cell labels, which is better replicated across different methods, than very fine-grained subtypes (work in preparation). Future studies could consider implementing additional approaches and rank cell types based on overlap between different methods. Additionally, in the approach of linking GWAS results to cell types, we leveraged the availability of single-cell RNAseq from non-diseased controls. It is plausible that in the diseased condition or genetic background, the same cell types in different brain regions may have distinguishable transcriptomic profiles. Future work could compare the predicted associated cell types obtained from a GWAS-to-cell-type framework versus from a framework that leverages single-cell RNAseq data from cases and controls.

## Acknowledgement

This project was supported by the NWO Gravitation grant BRAINSCAPES: a roadmap from neurogenetics to neurobiology (grant no. 024.004.012), the European Research Council Advanced Grant (grant no. ERC-2018-ADG 834057).

