## Supplementary Notes and Figures for "Identification of cell types associated with 14 brain phenotypes from more than 10 million single cells"

**Table of contents**

| Table of contents | Page |
| --- | --- |
| Supplementary Note 1. Detailed information on processing of snRNAseq data | 2 |
| Supplementary Note 2. GWAS selection criteria | 9 |
| Supplementary Note 3. Detailed method in identifying the main associated cell types | 10 |
| Supplementary Figure 1. Presence of broad cell type labels in curated snRNAseq studies | 11 |
| Supplementary Figure 2. Significantly associated cell types per phenotype for Cognitive processing speed | 12 |
| Supplementary Figure 3. Significantly associated cell types per phenotype for Well-being | 13 |
| Supplementary Figure 4. Significantly associated cell types per phenotype for Smoking initiation | 13 |
| Supplementary Figure 5. Significantly associated cell types per phenotype for MDD | 14 |
| Supplementary Figure 6. Significantly associated cell types per phenotype for SCZ | 14 |
| Supplementary Figure 7. Significantly associated cell types per phenotype for Alcohol consumption | 15 |
| Supplementary Figure 8. Significantly associated cell types per phenotype for Neuroticism | 15 |
| Supplementary Figure 9. Significantly associated cell types per phenotype for ADHD | 16 |
| Supplementary Figure 10. Significantly associated cell types per phenotype for Bipolar | 16 |
| Supplementary Figure 11. Significantly associated cell types per phenotype for Insomnia | 17 |
| Supplementary Figure 12. Significantly associated cell types per phenotype for Parkinson's | 17 |
| Supplementary Figure 13. Significantly associated cell types per phenotype for Migraine | 18 |
| Supplementary Figure 14. Significantly associated cell types per phenotype for Alzheimer's | 18 |
| Supplementary Figure 15. Significantly associated cell types per phenotype for BMI | 19 |

### Supplementary Note

#### Supplementary Note 1. Detailed information on processing of snRNAseq data

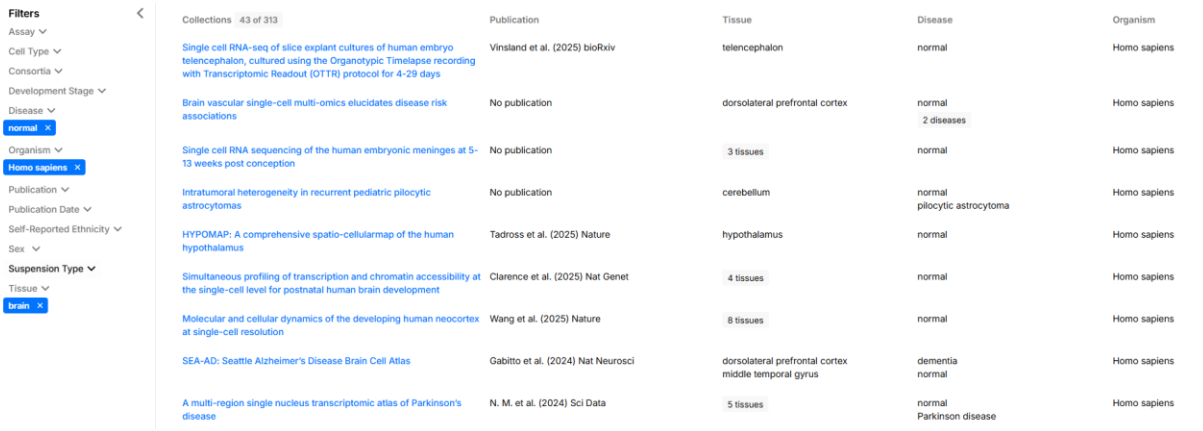


We downloaded the snRNAseq data in AnnData (h5ad) format from cellxgene. For each downloaded h5ad file, we analyzed the metadata (obtained from the "obs" layer of the anndata object) to obtain relevant information regarding developmental stages, brain regions, or disease status. We briefly described each dataset below. Additional detailed information can be found on the github page: https://github.com/tanyaphung/scrnaseq_viewer/tree/main/notes.

#### Datasets excluded

We excluded seven datasets from further processing: Chan et al. 2021 and Salcher et al. 2022 were excluded because these studies were focused on lung cancer. Tian et al. 2023 and Chien et al. 2023 were excluded because these data were methylation data and not gene expression data. He et al. 2024 and Vinsland et al. 2025 was excluded because this is an organoid dataset and we wanted to focus on human brain tissues in this study. Mannens et al. 2024 was excluded because this is a single-cell ATACseq dataset.

#### 1. Smith et al. 2021

Raw counts were downloaded from cellxgene (https://cellxgene.cziscience.com/collections/e02201d7-f49f-401f-baf0-1eb1406546c0) (access date: December 25 2023). Two files were downloaded: one for the midgestational human neocortex and one for the infant human neocortex. For the midgestational human cortex, there were 118,647 cells total that originated from samples at 22$^{nd}$ week post-fertilization human stage. The data were obtained from seven areas of the brain: parietal lobe, hippocampal formation, primary visual cortex, medial ganglionic eminence, caudal ganglionic eminence, orbitofrontal cortex, and anterior cingulate cortex.  For the infant human neocortex, there were 51,878 cells total that originated from samples at 7-month-old human stage. The data were obtained from four areas of the brain: prefrontal cortex, temporal lobe, parietal lobe, and primary visual cortex. We generated seven sub-datasets for the midgestational stage (Supplementary Table 2, ids 253-259) and four sub-datasets for the infant stage (Supplementary Table 2, ids 260-263).

2. Jakel et al. 2019

Raw counts were downloaded from cellxgene (https://cellxgene.cziscience.com/collections/16c1e722-96ae-4bf6-b408-cd7f8918484f) (access date: January 19 2024). The data originated from both non-diseased individuals and individuals with multiple sclerosis so we kept only the cells that originated from non-diseased individuals. The data were obtained from brain white matter and brain. The developmental stages indicated that these samples are at the late postnatal stage (ages were 35, 37, 44, 49, 57, 60, 64, and 82). There were 17,799 cells total in which 6,591 cells were from non-diseased samples.

3. Leng et al. 2021

Raw counts were downloaded from cellxgene (https://cellxgene.cziscience.com/collections/180bff9c-c8a5-4539-b13b-ddbc00d643e6) (access date: May 30 2024). The data originated from both non-diseased individuals and individuals with Alzheimer's Disease so we kept only the cells that originated from non-diseased individuals. The data were obtained from individuals at the late postnatal stage (ages were 50, 50, 71, 72, 77, 80, 82, and 87). There were 9,730 and 12,646 cells from the non-diseased entorhinal cortex and superior frontal gyrus, respectively. We generated two sub-datasets corresponding to two brain regions (Supplementary Table 3, ids 424 and 425).

4. Aldinger et al. 2021

Raw counts were downloaded from cellxgene (https://cellxgene.cziscience.com/collections/1b014f39-f202-45ae-bb7d-9286bddd8d8b) (access date: January 7 2024). The data originated from non-diseased individuals spanning nine developmental stages: 9, 10, 11, 12, 14, 17, 18, 20, and 21 week post-fertilization. There were 69,174 cells in total. We generated nine sub-datasets corresponding to the nine developmental stages (Supplementary Table 3, ids 264-272).

5. Bhaduri et al. 2021

Raw counts were downloaded from cellxgene (https://cellxgene.cziscience.com/collections/c8565c6a-01a1-435b-a549-f11b452a83a8) (access date: January 8 2024). The data originated from non-diseased individuals spanning across six brain regions: prefrontal cortex, primary motor cortex, parietal cortex, primary somatosensory cortex, temporal cortex, primary visual cortex, and across eight prenatal developmental stages: $14^{th}$ week, 16th, 17th, 18th, 19th, 20th, 22nd, and 25th week post-fertilization human stage. There were 457,965 cells in total. We generated 42 sub-datasets for each unique developmental stage and brain region combination (Supplementary Table 3, ids 276-317).

6. Bakken et al. 2021

Raw counts were downloaded from cellxgene (https://cellxgene.cziscience.com/collections/367d95c0-0eb0-4dae-8276-9407239421ee) (access date: January 8 2024). The data originated from non-diseased individuals from three organisms: Callithrix jacchus, Homo sapiens, Mus musculus. We kept only cells that originated from Homo sapiens. The data were sequenced from the primary motor cortex of late postnatal individuals (ages: 50 and 60). There were 20,490 total cells.

7. Kamath et al. 2022

Raw counts were downloaded from cellxgene (https://cellxgene.cziscience.com/collections/b0f0b447-ac37-45b0-b1bf-5c0b7d871120) (access date: May 30 2024). The data originated from non-diseased individuals and individuals with Lewy body dementia and Parkinson disease. We kept only cells from non-diseased individuals. The data were sequenced from the substantia nigra pars compacta from individuals at the late postnatal stage of development. In total there were 369,346 normal cells.

8. Otero-Garcia et al. 2022

Raw counts were downloaded from cellxgene (https://cellxgene.cziscience.com/collections/b953c942-f5d8-434f-9da7-e726ba7c1481) (access date: January 17 2024). The data originated from non-diseased individuals and individuals with Alzheimer's disease. We kept only cells from non-diseased individuals. The data were sequenced from the prefrontal cortex from individuals at the late postnatal stage of development. In total there were 57,534 normal cells.

9. Ma et al. 2022

Raw counts were downloaded from cellxgene (https://cellxgene.cziscience.com/collections/e1fa9900-3fc9-4b57-9dce-c95724c88716) (access date: January 16 2024). The data included cells from four organisms: Homo sapiens, Pan troglodytes, Macaca mulatta, Callithrix jacchus and only cells from Homo sapiens were kept The data were sequenced from the dorsolateral prefrontal cortex of non-diseased individuals at the late postnatal stages. In total there were 172,120 human cells.

10. Gabitto et al. 2023

Raw counts were downloaded from cellxgene (https://cellxgene.cziscience.com/collections/1ca90a2d-2943-483d-b678-b809bf464c30) (access date: June 12 2023). The data were sequenced from the middle temporal gyrus from non-diseased human individuals. There were a total of 164,053 cells.

11. Seeker et al. 2023

Raw counts were downloaded from cellxgene (https://cellxgene.cziscience.com/collections/9d63fcf1-5ca0-4006-8d8f-872f3327dbe9) (access date: January 16 2024). The data were sequenced from non-diseased individuals from three regions: Brodmann (1909) area 4, white matter of cerebellum, and cervical spinal cord white matter. The developmental stages were divided into young and old. We generated size sub-datasets corresponding to the developmental stages and brain region combinations. There were in total 45,528 cells.

12. Gittings et al. 2023

Raw counts were downloaded from cellxgene (https://cellxgene.cziscience.com/collections/aee9c366-f2fb-470b-8937-577d5d87d3fc) (access date: January 22 2024). The data originated from non-diseased individuals and individuals with amyotrophic lateral sclerosis and amyotrophic lateral sclerosis 26 with or without frontotemporal dementia. We kept only cells that originated from non-diseased individuals. The data were sequenced from the frontal cortex and the occipital cortex from individuals at the late postnatal stage. The samples were divided into two parts: C9-ALS, Cp-

ALS/FTD and C9-FTD. We generated four sub-datasets (Supplementary Table 3, ids 341-344). There were a total of 115,568 cells.

#### 13. Jorstad et al. 2023

Raw counts were downloaded from cellxgene (https://cellxgene.cziscience.com/collections/4dca242c-d302-4dba-a68f-4c61e7bad553) (access date: December 18 2023). The data were sequenced from non-diseased individuals at the late postnatal developmental stage from the middle temporal gyrus. There were a total of 156,285 cells. Since there were two different sequencing strategies, smartseq and 10x, we generated two sub-datasets corresponding to each sequencing technique (Supplementary Table 3, ids 226 and 227).

#### 14. Siletti et al. 2023

Raw counts were downloaded from cellxgene (https://cellxgene.cziscience.com/collections/283d65eb-dd53-496d-adb7-7570c7caa443) (access date: May 04 2023). The data were sequenced from non-diseased individuals at the late postnatal developmental stage from ten brain regions and from multiple dissections within each region. There were a total of 3,369,399 cells. We generated 105 sub-datasets corresponding to each dissection (Supplementary Table 3, ids 7-111).

#### 15. Zhu et al. 2023

Raw counts were downloaded from cellxgene (https://cellxgene.cziscience.com/collections/ceb895f4-ff9f-403a-b7c3-187a9657ac2c) (access date: December 24 2023). The data were sequenced from non-diseased individuals from the cortical plate and the dorsolateral prefrontal cortex. There were six stages of development: early fetal, late fetal, infancy, childhood, adolescence, and adulthood. There were a total of 45,549 cells. We generated six sub-datasets for each unique brain region and developmental stage combination (Supplementary Table 3, ids 247-252).

#### 16. Jorstad et al. 2023

Raw counts were downloaded from cellxgene (https://cellxgene.cziscience.com/collections/d17249d2-0e6e-4500-abb8-e6c93fa1ac6f) (access date: January 10 2024). The data were sequenced from non-diseased individuals at the late postnatal developmental stages from these brain regions: Primary motor cortex (M1), Primary somatosensory cortex (S1), Primary auditory cortex (A1), Primary visual cortex (V1), Dorsolateral prefrontal cortex (DFC), Anterior cingulate cortex (ACC), Middle temporal gyrus (MTG), and Angular gyrus (AnG). There were 1,155,822 cells that were sequenced from 10x technology and 49,417 cells that were sequenced from Smart-seq technology. We generated 8 and 6 sub-datasets corresponding to the brain region for each sequencing technology (Supplementary Table 3, ids 318-331).

#### 17. Velmeshev et al. 2023

Raw counts were downloaded from cellxgene (https://cellxgene.cziscience.com/collections/bacccb91-066d-4453-b70e-59de0b4598cd) (access date: December 19 2023). Data originated from samples from second trimester to adulthood and from ganglionic eminences and from the cortex (prefrontal, cingulate, temporal, insular, and motor cortices). There were 413,682 nuclei originally and 358,663 nuclei after filtering. After integrating with existing datasets, there were 709,372 nuclei in this dataset. We generated 58 sub-datasets for each developmental stage and brain region combination (Supplementary Table 3, ids 366-423).

#### 18. Sepp et al. 2023

Raw counts were downloaded from cellxgene (https://cellxgene.cziscience.com/collections/72d37bc9-76cc-442d-9131-da0e273862db) (access date: December 21 2023). Data were sequenced from the cerebellum across developmental stages: early neurogenesis to adulthood: 7 week post conception (wpc), 8 wpc, 9 wpc, 11 wpc, 17 wpc, 20 wpc, newborn infant, toddler, and adult. There were 163,283 cells from non-diseased individuals. We generated 11 sub-datasets corresponding to the different developmental stages (Supplementary Table 3, ids 236-243).

#### 19. Phan et al. 2024

Raw counts were downloaded from cellxgene (https://cellxgene.cziscience.com/collections/cec4ef8e-1e70-49a2-ae43-1e6bf1fd5978) (access date: May 30 2024). Data originated from non-diseased individuals and individuals with opiate dependence. Thus, only cells from non-diseased individuals were kept. Data were sequenced from the caudate nucleus and putamen from late postnatal individuals. In total, there were 44,449 cells in this dataset. We generated two sub-datasets corresponding to the two regions (Supplementary Table 3, ids 427-428).

#### 20. Nascimento et al.

Raw counts were downloaded from cellxgene (https://cellxgene.cziscience.com/collections/cae8bad0-39e9-4771-85a7-822b0e06de9f) (access date: January 23 2024). Data originated from non-diseased individuals from the entorhinal cortex and ganglionic eminence. Data spans different developmental timepoint: Fetal, Infant, Toddler, Teen, and Adult. In total, there were 153,005 cells in this dataset. We generated 21 sub-datasets for each brain region and developmental stage combination (Supplementary Table 3, ids 345-365).

#### 21. TorresFlores et al.

Raw counts were downloaded from cellxgene (https://cellxgene.cziscience.com/collections/10bf5c50-8d85-4c5f-94b4-22c1363d9f31) (access date: January 10 2024). Data originated from non-diseased individuals and individuals with pilocytic astrocytoma. Therefore, only cells from non-diseased individuals were kept. Data were sequenced from the cerebellum from individuals at the child developmental stage. There were 1,346 cells in this dataset.

#### 22. Wang et al. 2024

Raw counts were downloaded from cellxgene (https://cellxgene.cziscience.com/collections/ad2149fc-19c5-41de-8cfe-44710fbada73) (access date: February 9 2025). Data originated from five developmental stages (first trimester, second trimester, third trimester, infancy, and adolescence) and eight unique brain regions (neocortex, telencephalon, forebrain, prefrontal cortex, visual cortex, brodmann area 10, 17, and 9). Note that at each developmental stage, not all of the eight brain regions were sampled; thus we generated 14 sub-datasets for each unique developmental stage and brain region combination (Supplementary Table 3, internal ids 429-442). There are 232,327 cells total in this dataset.

23. Braun et al. 2023

Raw counts were downloaded from cellxgene (https://cellxgene.cziscience.com/collections/4d8fed08-2d6d-4692-b5ea-464f1d072077) (access date: February 10 2025). Data originated from twelve stages during the first trimester: Carnegie stage 15, 16, 18, 20, 22, and 9, 10, 11, 12, 13, 14, 15 weeks post fertilization. There were ten brain regions: brain, cerebellum, diencephalon, forebrain, head, hindbrain, medulla, midbrain, pons, and telencephalon. We generated 50 sub-datasets for each unique developmental age and brain region combination (Supplementary Table 3, internal ids 443-492). There are approximately 1.6 million cells in this dataset.

24. Rexach et al. 2024

Raw counts were downloaded from cellxgene (https://cellxgene.cziscience.com/collections/c53573b2-eff4-4c5e-9ad0-b24d422dfd9b) (access date: February 11 2025). Data originated from adult human brain from three brain regions: primary visual cortex, insular cortex, and brodmann area 4. Only cells from non-diseased normal samples were included (98,794 cells). We generated three sub-datasets for each brain region (Supplementary Table 3, internal ids 494-496).

25. Dharshini et al. 2024

Raw counts were downloaded from cellxgene (https://cellxgene.cziscience.com/collections/0d35c0fd-ef0b-4b70-bce6-645a4660e5fa) (access date: February 11 2025). Data originated from adult human brain from three brain regions: brodmann area 7, 9, and 17. Only cells from non-diseased normal samples were included (113,755 cells). We generated three sub-datasets for each brain region (Supplementary Table 3, internal ids 497-499).

26. N.M. et al. 2024

Raw counts were downloaded from cellxgene (https://cellxgene.cziscience.com/collections/d5d0df8f-4eee-49d8-a221-a288f50a1590) (access date: February 12 2025). Data originated from adult human brain from five regions: dorsal motor nucleus of vagus nerve, primary visual cortex, prefrontal cortex, primary motor cortex, and medial globus pallidus. Only cells from non-diseased normal samples were included (445,297 cells). We generated five sub-datasets for each brain region (Supplementary Table 3, internal ids 500-504).

27. Clarence et al. 2025

Raw counts were downloaded from cellxgene (https://cellxgene.cziscience.com/collections/f406a653-c079-4bf9-aab6-85846c27571d) (access date: February 12 2025). Data originated from five developmental ages: infant stage, 4, 6, 14, 20, 39, 61, and 62 year old stage. We grouped the developmental ages into two developmental groups: the postnatal early group consist of the infant stage, 4, and 6 year old stages and the postnatal late group consist of 14, 20, 39, 61, and 62 year old stage. There were four tissues: caudate nucleus, hippocampal formation, dorsolateral prefrontal cortex, and anterior cingulate cortex. We generated nine sub-datasets for each developmental  group and brain region combination (Supplementary Table 3, internal ids 505-512). In total there are 101,924 cells.

#### 28. Tadross et al. 2025

Raw counts were downloaded from cellxgene (https://cellxgene.cziscience.com/collections/d0941303-7ce3-4422-9249-cf31eb98c480) (access date: February 12 2025). Data originated from twenty regions of the hypothalamus (note that while there were 21 unique values in the tissue column in the metadata, the labels Thalamaus was most likely mis-spelled; therefore, we grouped cells with cell type annotation Thalamus and Thalamaus to be from the thalamus). We generated twenty sub-datasets for each region (Supplementary Table 3, internal ids 514-533). In total there are 433,369 cells.

#### 29. Pan et al. 2024

Raw counts were downloaded from cellxgene (https://cellxgene.cziscience.com/collections/d0941303-7ce3-4422-9249-cf31eb98c480) (access date: February 13 2025). When considering only cells from the non-diseased normal samples, these cells originated from  the prefrontal cortex and there are 31,971 cells. We generated one sub-dataset (Supplementary Table 3, internal id 513).

### 30. Xu et al. 2023

Raw counts were downloaded from cellxgene (https://cellxgene.cziscience.com/collections/854c0855-23ad-4362-8b77-6b1639e7a9fc) (access date: March 12 2025). While the dataset included multiple tissues, we selected only the brain, specifically hippocampal formation. Xu et al. 2023 developed CellHint, a method to harmonize cell type annotation from different studies. As such, the dataset for hippocampal formation came from four studies: Ayhan et al. 2021, Franjic et al. 2022, Tran et al. 2021, and Siletti et al. 2022. We generated four sub-datasets corresponding to each dataset (Supplementary Table 3, internal ids 534-537).

#### 31. Johansen et al. 2023

Raw counts were downloaded from cellxgene (https://cellxgene.cziscience.com/collections/35928d1c-36fc-4f93-9a8d-0b921ab41745) (access date: March 12 2025). Data originated from the neocortex of adult human brain.

#### 32. Han et al. 2020

Raw counts were downloaded from cellxgene (https://cellxgene.cziscience.com/collections/38833785-fac5-48fd-944a-0f62a4c23ed1) (access date: September 2 2025). We subsetted to cells from the Adult Cerebellum and Fetal Brain only.

#### 33. Cao et al. 2020

Raw counts were downloaded from cellxgene (https://cellxgene.cziscience.com/collections/c114c20f-1ef4-49a5-9c2e-d965787fb90c

) (access date: September 3 2025). We subsetted to cells from the cerebellum and telencephalon only.

#### 34. Linnarsson

This is a dataset sequenced from the fetal human meninges. Raw counts were downloaded from cellxgene (https://cellxgene.cziscience.com/collections/7d66d871-091f-4602-9f42-85f86d2853e0) (access date: September 3 2025).

#### 35. Yang

This is a dataset sequenced from the dorsolateral prefrontal cortex. Raw counts were downloaded from cellxgene (https://cellxgene.cziscience.com/collections/433700dc-e8a5-48b0-b5cd-beb22f3f88fe) (access date: September 3 2025). We subsetted the data to cells from normal non-diseased samples only.

#### 36. Lee et al. 2024

To reduce computational burden, we only processed the RADC cohort from this dataset. Raw counts for the RADC cohort were downloaded from cellxgene (https://cellxgene.cziscience.com/collections/84ce6837-548d-4a1f-919f-0bc0d9a3952f) (access date: September 2 2025). We subsetted the data to cells from normal non-diseased samples only.

### Supplementary Note 2. GWAS selection criteria

For similar phenotypes, we selected a GWAS with highest sample size and/or highest proportion of cases. Between cognitive processing speed (Li et al. 2024, sample size 2,266,733) and cognitive processing accuracy (Li et al. 2024, sample size 1,783,727), we selected cognitive processing speed. Between smoking initiation (Liu et al. 2019, sample size 1,232,091), smoking cessation (Liu et al. 2019, sample size 547,219), and smoking status (Linner et al. 2019, sample size 518633), we selected smoking initiation. While the seven anxiety phenotypes with total sample size greater than 500,000 individuals from Brasher et al. 2023, we decided to exclude this phenotype due to very low proportion of cases (anxiety GWAS with the highest proportion of cases is around 4 percent). Across phenotypes representing depression, we selected the summary statistics from a meta-analysis of approximately 4 million European individuals (Adams et al. 2025). For schizophrenia, we selected the summary statistics from the Psychiatric Genomics Consortium due to higher proportion of cases. For Parkinson's disease, even though the GWAS from Kim et al. 2024 contains the largest number of cases (49049 cases and 18785 proxy cases), the GWAS summary statistics is not publicly available, we selected the GWAS from Nalls et al. 2021 which included approximately 17675 cases and 239152 controls. For migraine, we selected the summary statistics from Choquet et al. 2021 because the GWAS from Hautakangas with more migraine cases is not available publicly. For Alzheimer's disease, we selected the summary statistics for non 23andme individuals from the PGC-ALZ3 GWAS.

### Supplementary Note 3. Detailed method in identifying the main associated cell types

The following dendograms are for illustrative purpose of depicting the clusters of cell types after step 3.

#### Smoking initiation

##### Prenatal

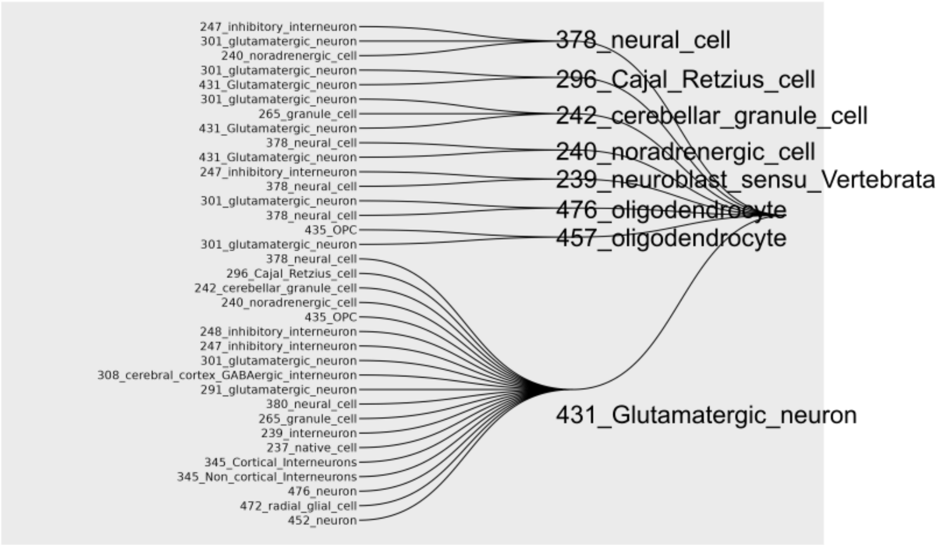


##### Postnatal

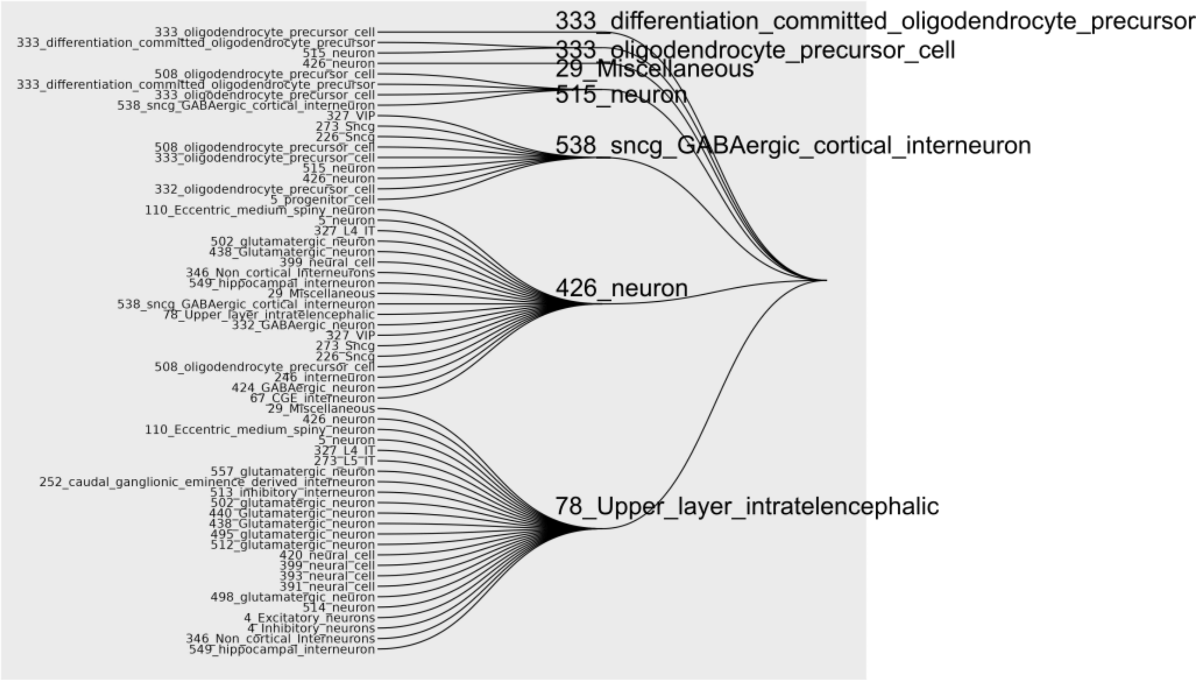


In each dendrogram, each node is the main cell type and the leaves are the cell types that were conditioned on. What is plotted are the main cell type – conditional cell types combinations that were forward selected (see Supplementary Table 1 from Watanabe et al. 2019). Then, each cluster is represented by the main cell type, which is represented by squares in Figure 3 and 4.

### Supplementary Figures

**Supplementary Figure 1. Presence of broad cell type labels in curated snRNAseq studies.**
Top panel is for level 1 that consists of progenitor, neuron, glia, and other. Bottom panel is for level 2. Each point is the proportion of the cell type computed within each dataset.

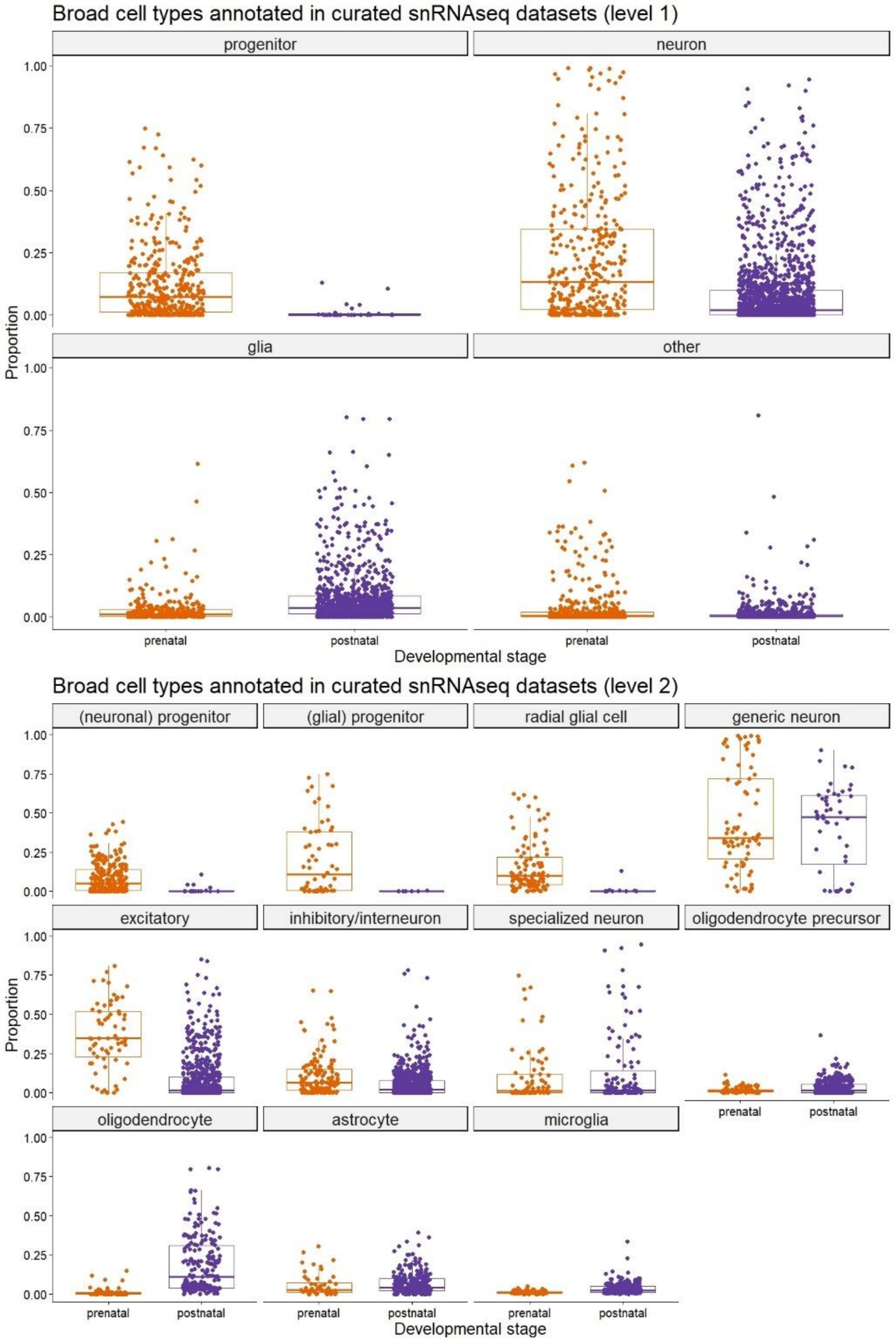

Supplementary Figure 2. Significantly associated cell types per phenotype for Cognitive processing speed

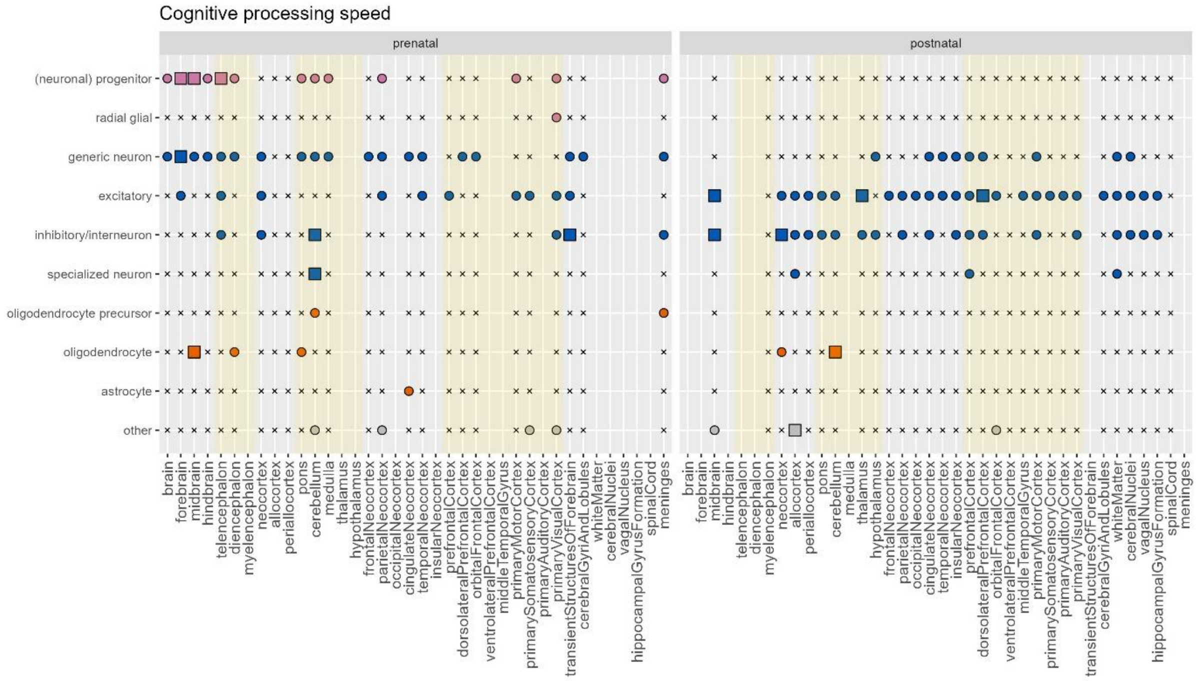

### Supplementary Figure 3. Significantly associated cell types per phenotype for Well-being

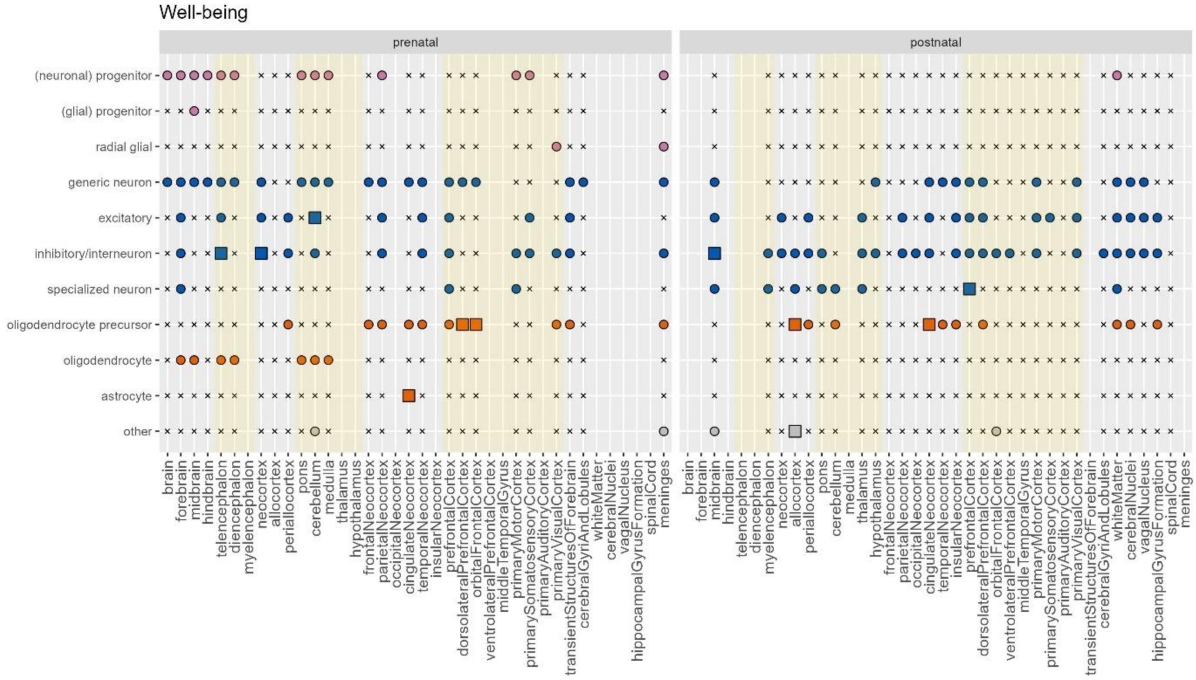


### Supplementary Figure 4. Significantly associated cell types per phenotype for Smoking initiation

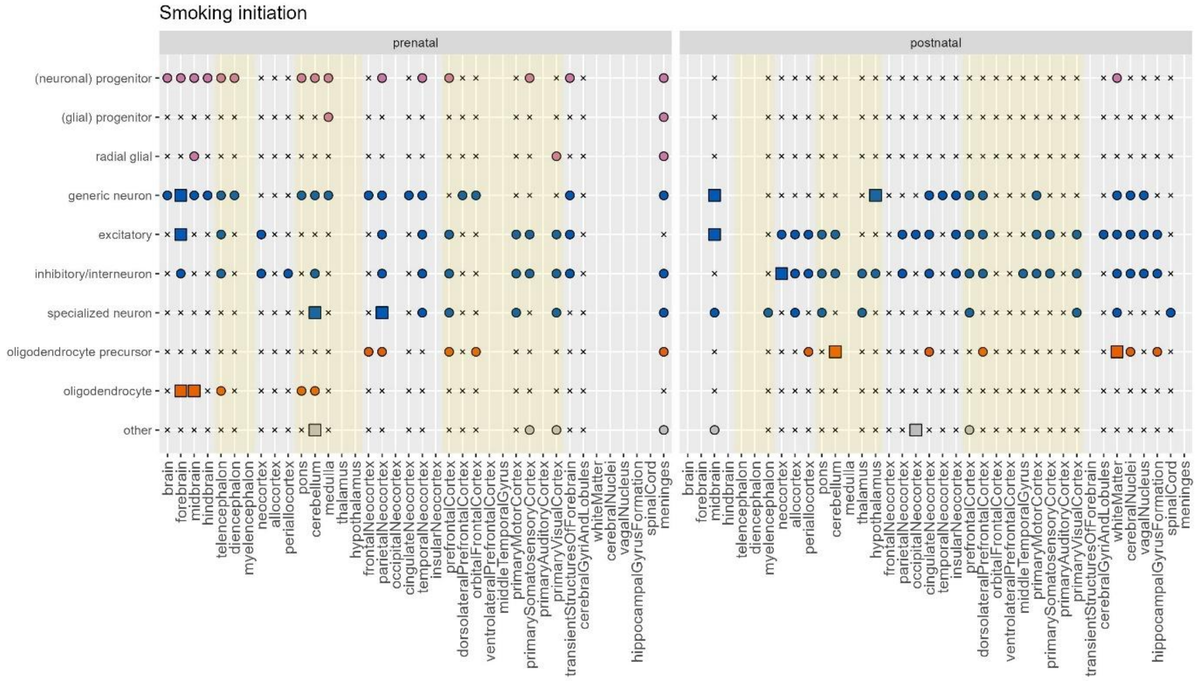

Supplementary Figure 5. Significantly associated cell types per phenotype for MDD

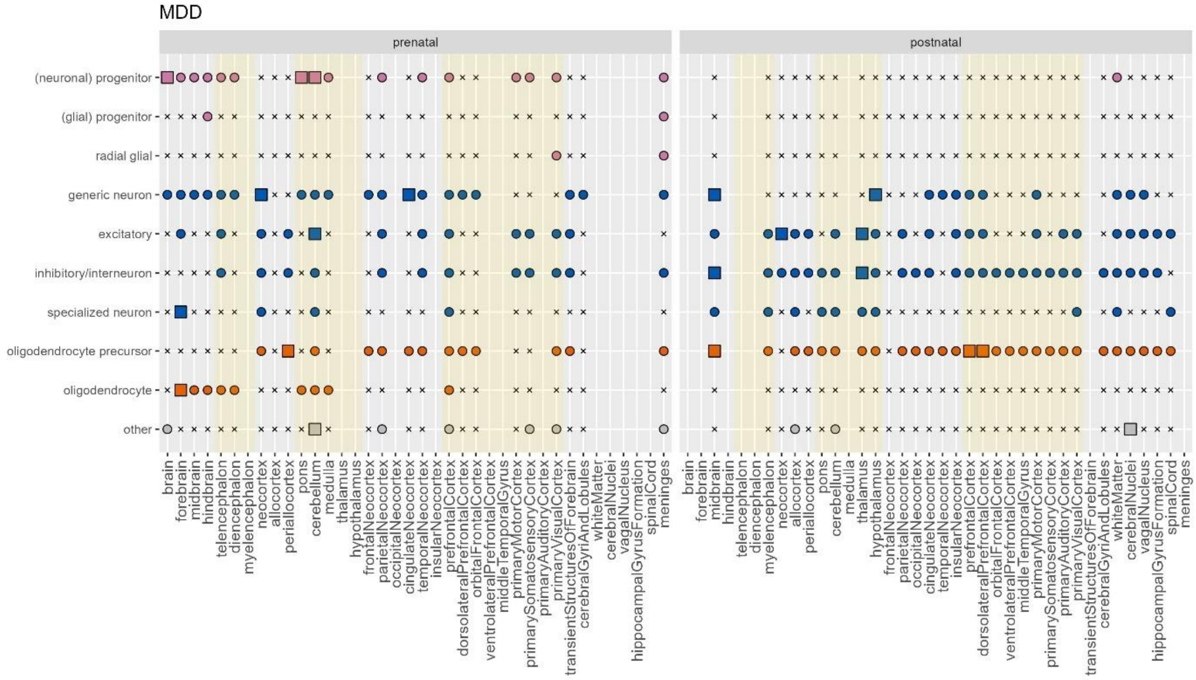


Supplementary Figure 6. Significantly associated cell types per phenotype for SCZ

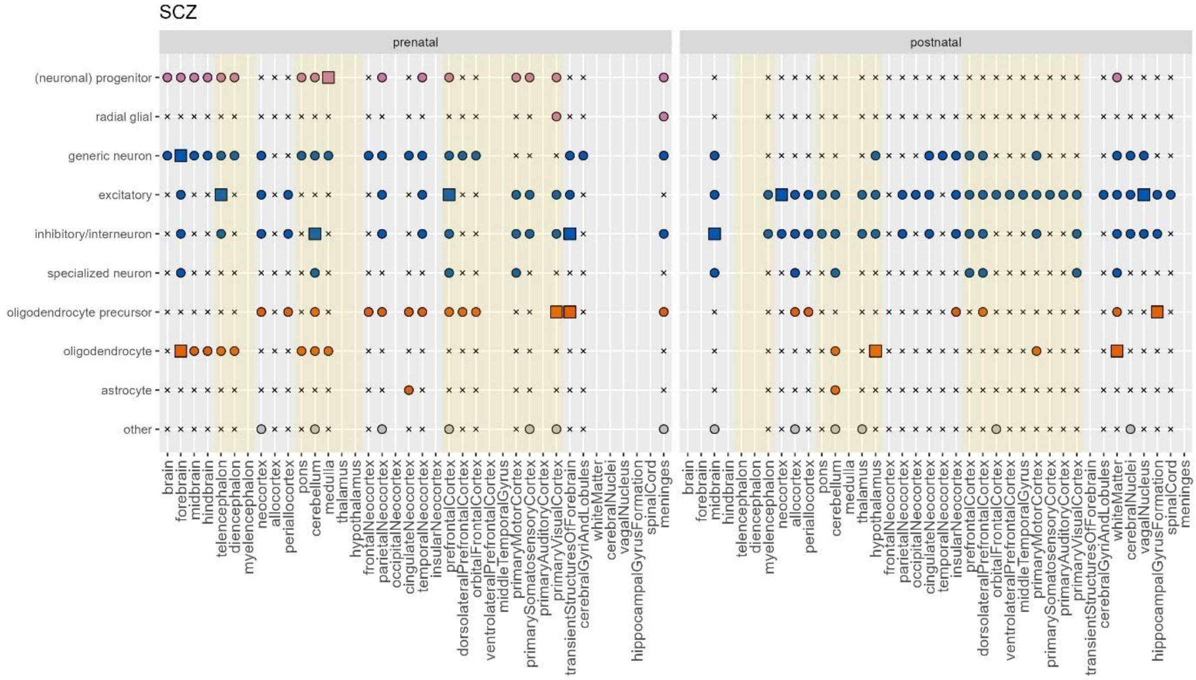

### Supplementary Figure 7. Significantly associated cell types per phenotype for Alcohol consumption

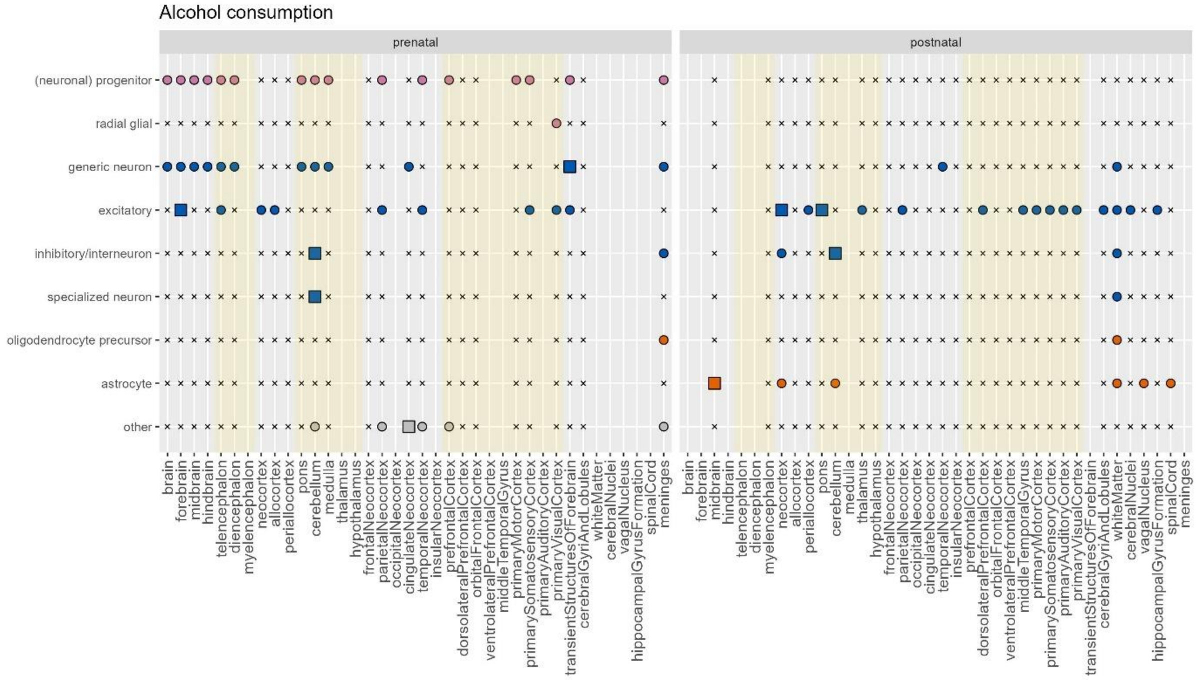


### Supplementary Figure 8. Significantly associated cell types per phenotype for Neuroticism

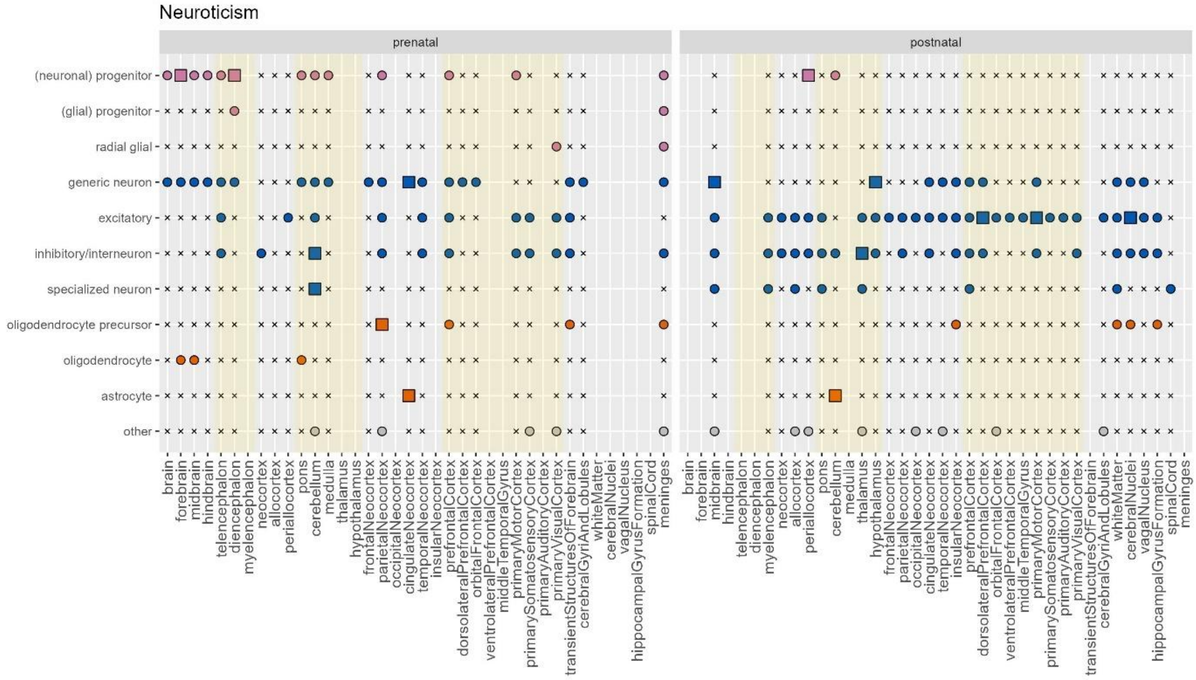

### Supplementary Figure 9. Significantly associated cell types per phenotype for ADHD

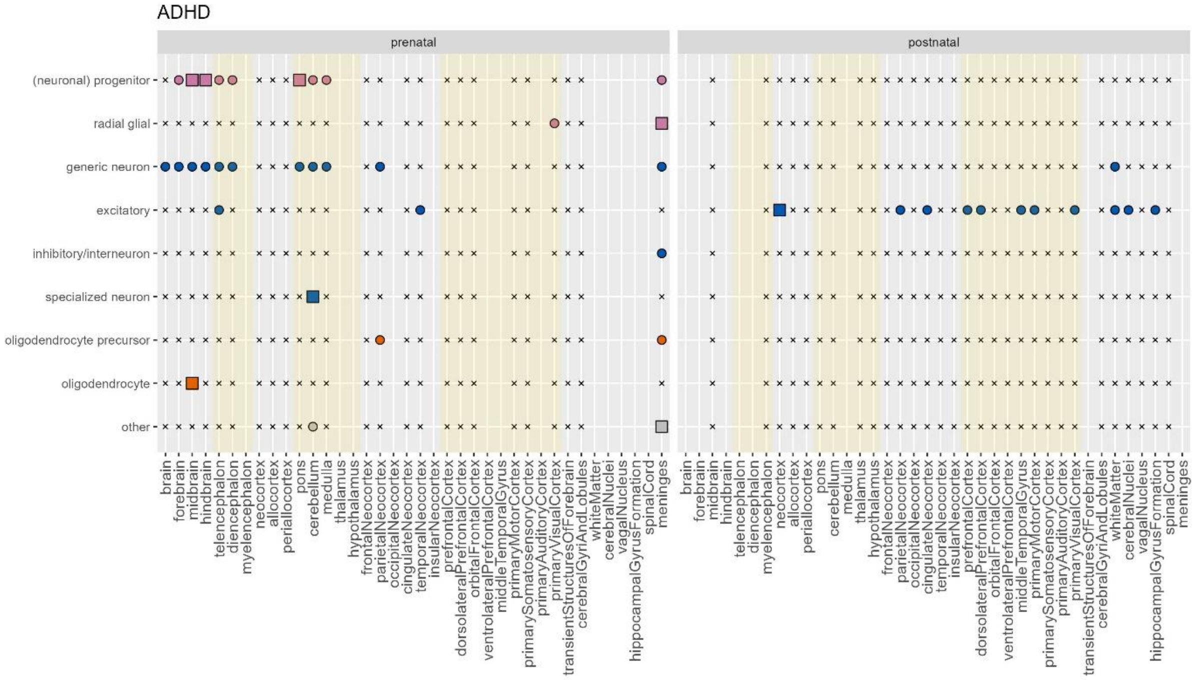


### Supplementary Figure 10. Significantly associated cell types per phenotype for Bipolar

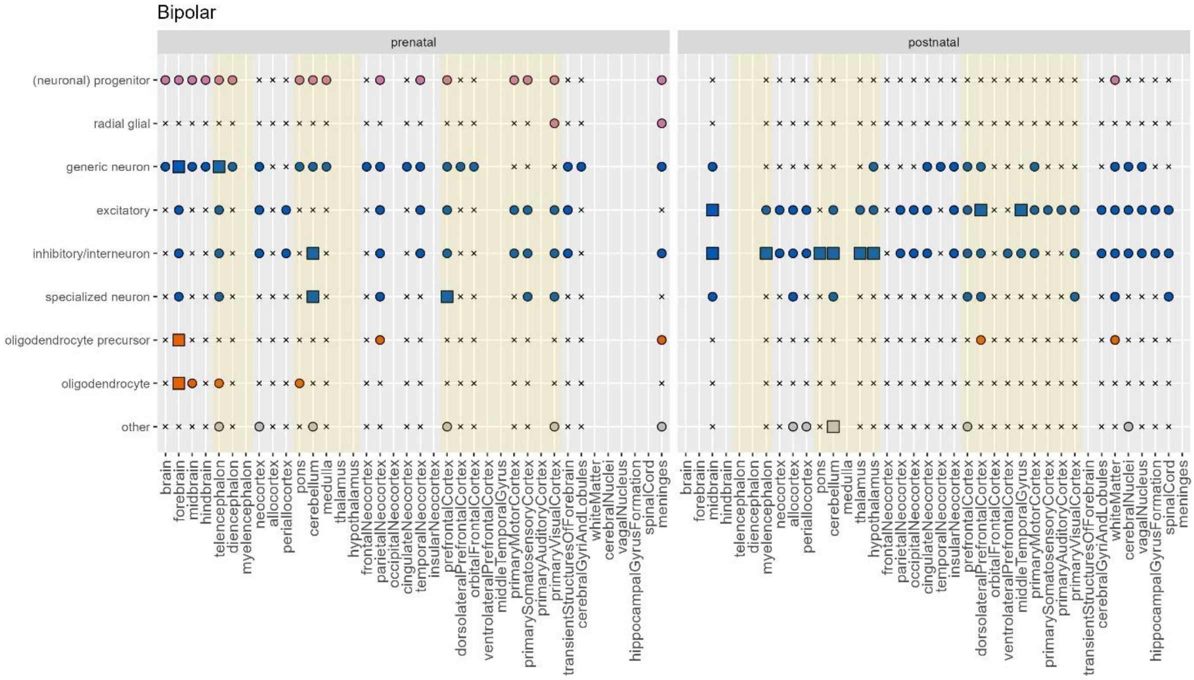

### Supplementary Figure 11. Significantly associated cell types per phenotype for Insomnia

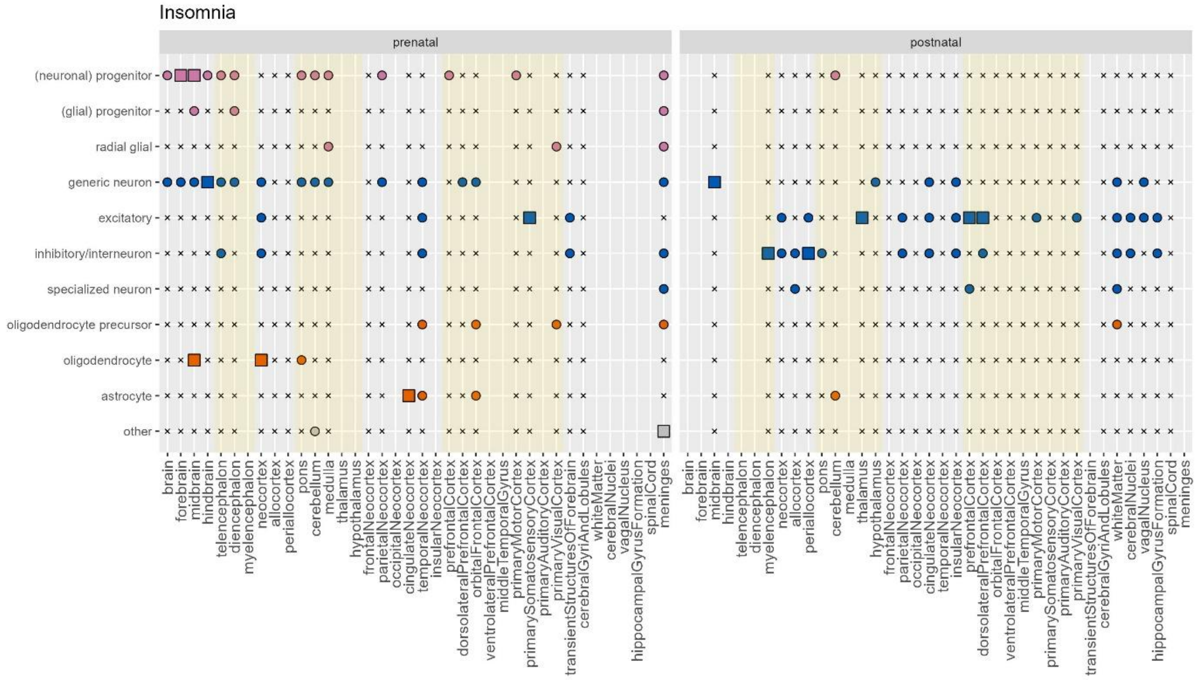


### Supplementary Figure 12. Significantly associated cell types per phenotype for Parkinson's

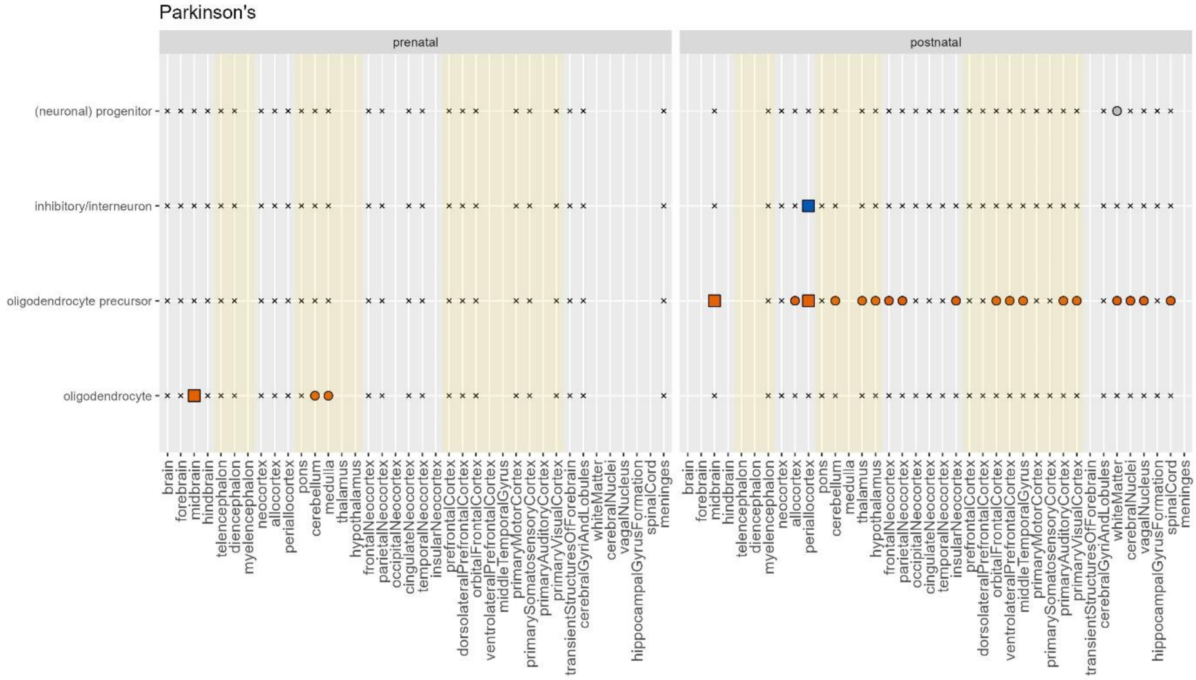

Supplementary Figure 13. Significantly associated cell types per phenotype for Migraine

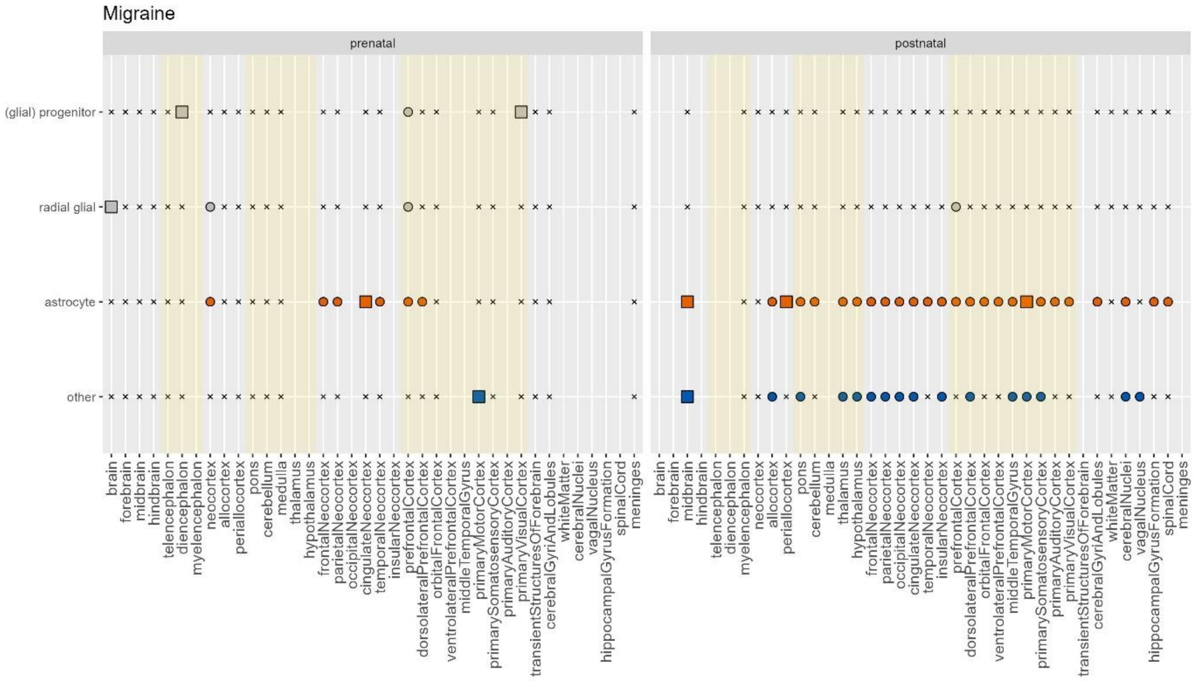


Supplementary Figure 14. Significantly associated cell types per phenotype for Alzheimer's

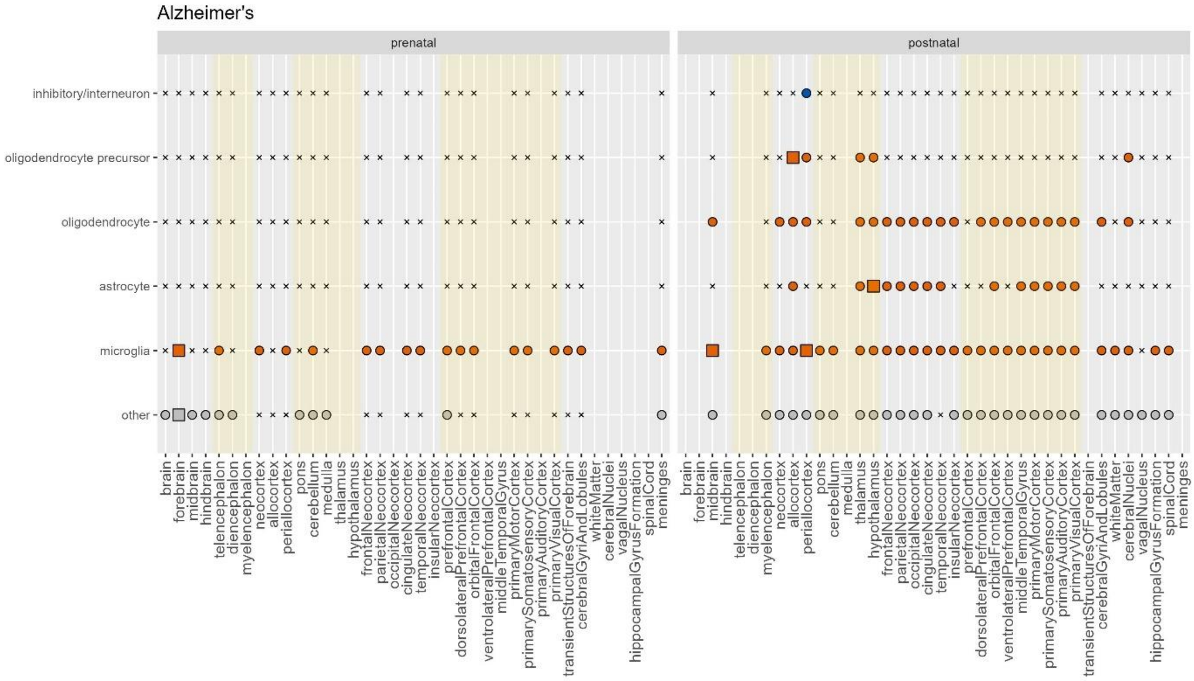

Supplementary Figure 15. Significantly associated cell types per phenotype for BMI

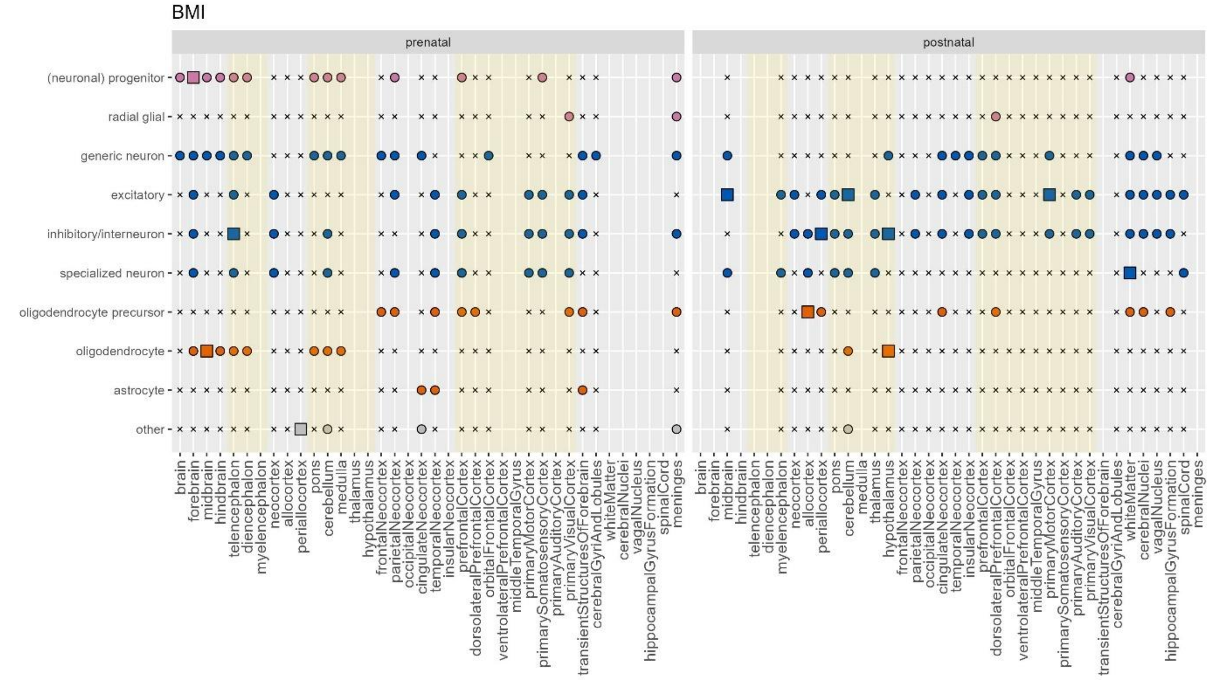
